# Stress induces DCP2 translation via a stalling-dependent mechanism

**DOI:** 10.64898/2026.08.25.746704

**Authors:** Roiuk Mykola, Marilena Neff, Taja Vatovec, Aram Papazian, Volkhard Helms, Aurelio A. Teleman

**Affiliations:** German Cancer Research Center (DKFZ) Heidelberg, Division B140, 69120 Heidelberg, Germany; Faculty of Medicine, Heidelberg University, 69120 Heidelberg, Germany; Faculty of Biosciences, Heidelberg University, 69120 Heidelberg, Germany; Max Planck Institute for Brain Research, Frankfurt, Germany; Gene Center and Department of Biochemistry, Ludwig- Maximilians-Universität, Munich, Germany; Center for Bioinformatics, Saarland University, Saarland Informatics Campus, Saarbrücken, Saarland, Germany

## Abstract

The integrated stress response (ISR) globally suppresses protein synthesis while selectively permitting translation of a small subset of stress-responsive mRNAs, many of which contain upstream or overlapping open reading frames (uORFs/oORFs). Although translational induction of transcripts such as ATF4 has classically been attributed to delayed re-initiation caused by reduced ternary complex availability, the mechanisms by which uORFs and oORFs allow ISR-selective translation remain incompletely understood. Here, using ribosome profiling during early ISR activation combined with reporter assays, we identify DCP2, encoding a major mRNA decapping enzyme, as a previously unrecognized ISR-induced transcript. We show that translational induction of DCP2 depends on an overlapping ORF whose conserved 3′ region, corresponding to a ribosome pausing site, acts as a potent inhibitory element. Both the DCP2 oORF and main ORF increase in translation during stress, indicating that stress relieves repression by this inhibitory element. This reveals a mode of ISR-dependent gene regulation in which inhibition by a nascent peptide or stalling element embedded either in a uORF or an oORF is relieved upon stress to induce translation.

## Introduction

Cells must constantly adapt to fluctuating environments. Because mRNA translation represents the final step in the gene expression program, and is the closest step to the protein product, it provides a rapid and tunable mechanism for adjusting cellular physiology. Importantly, translation is not a simple one-to-one conversion of mRNA into protein; rather, cells can selectively modulate the translation of specific subsets of mRNAs, controlling the amount of protein produced from each transcript.

One pervasive type of environmental condition is stress, such as hypoxia or nutrient deprivation. Such stresses are especially relevant in cancer, where cells in poorly vascularized tumor regions face chronic nutrient and oxygen limitation. Cells sense stress primarily through two central pathways: mTORC1 and the integrated stress response (ISR). Both pathways reduce global protein synthesis, conserving energy and resources under unfavorable conditions (Wek & Staschke, 2010). A plethora of stresses inactivate mTORC1, reducing phosphorylation of its target 4E-BP, which then binds eIF4E and limits ribosome recruitment to mRNAs (Heberle *et al*, 2015; Proud, 2019) and hence global protein synthesis. Similarly, the ISR (Costa-Mattioli & Walter, 2020; Pakos-Zebrucka *et al*, 2016) detects a variety of stresses such as amino acid limitation, hypoxia, or ER stress through four kinases that converge on the alpha subunit of eIF2 (eIF2α). Phosphorylation of eIF2α increases its affinity for the guanine exchange factor eIF2B, acting as a competitive inhibitor (Adomavicius *et al*, 2019; Bogorad *et al*, 2017; Kashiwagi *et al*, 2019; Kenner *et al*, 2019; Kwon *et al*, 2017). This reduces cytosolic eIF2-GTP and ternary complex levels, thereby suppressing initiator tRNA recruitment to ribosomes and global translation.

This global reduction in protein synthesis is energetically sensible: translation is one of the most resource-intensive processes in the cell, consuming both energy and amino acids. Yet cells cannot simply halt translation - they must respond to stress in order to survive. Accordingly, a subset of stress-response mRNAs evades this translational repression, such as ATF4, ATF5, and DDIT3 (Andreev *et al*, 2015; Reich *et al*, 2020; Sidrauski *et al*, 2015). The mechanisms enabling certain transcripts to evade repression remains incompletely understood.

A common feature of these resistant transcripts is the presence of upstream or overlapping open reading frames (uORFs and oORFs) in the 5’ untranslated region (5’UTR) (Andreev *et al.*, 2015; Johnstone *et al*, 2016). In this manuscript, we use the term uORF to denote short coding sequences that start and stop in the 5’UTR (upstream), and oORFs to denote open reading frames that start in the 5’UTR but extend beyond the main ORF in a different reading frame (overlapping). Typically, uORFs and oORFs are inhibitory: uORFs can sequester the scanning 43S pre-initiation complex, reducing initiation at the downstream AUG. Cells can bypass this repression via leaky scanning (Bohlen *et al*, 2023; Dever *et al*, 2023; Smirnova *et al*, 2022; Wang & Rothnagel, 2004; Weiss & Dikstein, 2024) or translation reinitiation (Skabkin *et al*, 2013), whereby the ribosome translates a uORF and then resumes scanning for a downstream AUG. In the case of oORFs, reinitiation is ineffective because translation terminates downstream of the main ORF (mORF) start codon and scanning is predominantly unidirectional (Pestova & Kolupaeva, 2002), so the only way to bypass an oORF is through leaky scanning. Although uORFs and oORFs are generally suppressive under basal conditions, they can paradoxically enable selective translation of stress-responsive transcripts (Dever *et al.*, 2023). The precise mechanisms remain an active area of investigation.

A canonical example of a stress-responsive transcript is ATF4 (Harding *et al*, 2000; Vattem & Wek, 2004), a transcription factor that drives expression of numerous stress-response genes. ATF4 contains two uORFs (one of which is a start codon followed immediately by a stop codon) and one oORF that regulate its translation upon stress. Its activation has traditionally been explained by delayed reinitiation (REI) (Lu *et al*, 2004; Vattem & Wek, 2004). In this model, the ribosome translates the two uORFs, and then continues scanning to reinitiate further downstream. Under normal conditions, abundant ternary complex activity promotes rapid delivery of a new initiator tRNA to the scanning 43S, thus enabling recognition of the oORF AUG and suppression of translation of ATF4’s mORF. During ISR activation, ternary complex levels are reduced, delaying reinitiation and allowing the ribosome to bypass the oORF AUG, initiating instead at the downstream main ORF. In support of this model, manipulations that increase the length of the interspace between the end of the second uORF and the oORF cause suppression of ATF4 activation (Vattem & Wek, 2004). However, recent studies challenge aspects of this model. Ribosome profiling and oORF peptide detection indicate that oORF translation increases, rather than decreases, upon ISR activation (Andreev *et al.*, 2015; Sidrauski *et al.*, 2015; Starck *et al*, 2016; Zhou *et al*, 2018). A recent, thorough study by the Valasek lab found that multiple regulatory mechanisms act additively to yield full ATF4 activation, and that part of the mechanism involves a stem-loop in the ATF4 oORF as well as mRNA methylation (Smirnova *et al*, 2024).

Understanding how cells control translation in response to stress is not only of fundamental interest, but also has broad implications for cancer and neurodegenerative diseases. We previously studied how cells employ alternative initiation mechanisms when mTORC1 is inhibited and eIF4E activity is impaired (Roiuk *et al*, 2024). Here, we focus on elucidating how cells regulate translation when the other major stress-response pathway, the ISR, is active.

## Results

### Identification of an early timepoint upon activation of the integrated stress response with translational reprogramming but few transcriptional changes

To study the translational changes that happen upon activation of the integrated stress response (ISR), we aimed to activate the ISR and to perform ribosome profiling. We employed tunicamycin, an antibiotic that primarily inhibits N-linked glycosylation, thereby inducing the unfolded protein response (UPR) in the endoplasmic reticulum. The UPR consists of three signalling branches (Hetz *et al*, 2020): one that regulates translation through PERK-mediated phosphorylation of eIF2α and two that cause transcriptional reprogramming. Hence, we aimed to identify an early timepoint after tunicamycin treatment where translational effects are visible and robust, but transcriptional effects are still minimal. To this end, we monitored the phosphorylation of eIF2α and the accumulation of ATF4, a downstream target of the ISR, and found that both are increased at 2 hours of treatment (Suppl. Fig. 1A-C). Consistent with this, polysome profiling revealed a strong drop in the polysome to monosome ratio at 2 hours, indicating global repression of translation (Suppl. Fig. 1D). Although this 2-hour time point already shows some activation of the IRE1-mediated transcriptional branch of the UPR, as indicated by accumulation of the spliced form of Xbp1 (Suppl. Fig. 1E), these effects are less pronounced than at later time points. Of note, longer treatments with tunicamycin, which we studied previously (16 hours) (Roiuk *et al*, 2025), additionally cause suppression of the mTORC1 pathway (Suppl. Fig. 1F), likely due in part to upregulation of DDIT4 and Sestrin 2 (SESN2), as previously reported (Condon *et al*, 2021; Figlia *et al*, 2025; Whitney *et al*, 2009; Ye *et al*, 2015) (and see below). Therefore, we decided to perform ribosome profiling at 2 hours after tunicamycin treatment, when translation is actively reprogrammed but the transcriptional impact is still mild and can be taken into account by normalizing ribosome footprints to total mRNA levels.

Analysis of our ribosome profiling data revealed enrichment of footprints in coding sequences as well as the expected triplet periodicity (Suppl. Fig. 2A), attesting to the quality of the data. Interestingly, tunicamycin treatment caused increased occupancy of 80S ribosomes on start codons, suggesting longer pausing for unknown reasons which might be worth studying in the future (Suppl. Fig. 2A). Overall, our ribosome profiling data identified a relatively small subset of transcripts that were affected, predominantly at the level of translation, as expected by the early timepoint we selected, and consistent with previous reports (Andreev *et al.*, 2015; Sidrauski *et al.*, 2015) (Fig. 1A, Table S1). Most responsive transcripts exhibited translational reprogramming with minimal changes in mRNA abundance, including canonical ISR targets such as ATF4, IFRD, and PPP1R15A (Fig. 1A). We also observed transcriptional upregulation of *HERPUD, HSPA5, DNAJN8, ATF3*, and *MIS12*. With the exception of *MIS12*, these genes have previously been shown to be transcriptionally induced during the unfolded protein response (UPR) (Hashimoto *et al*, 2002; Heldens *et al*, 2011; Ho & Chan, 2015; Yoshida *et al*, 2001) (Fig. 1A). In contrast to this 2-hour time point, reanalysis of data we previously generated at 16- hours of tunicamycin treatment revealed extensive transcriptional and translational remodeling, including pronounced upregulation of SESN2 and DDIT4 (Suppl. Fig. 2C).

**Figure 1.**
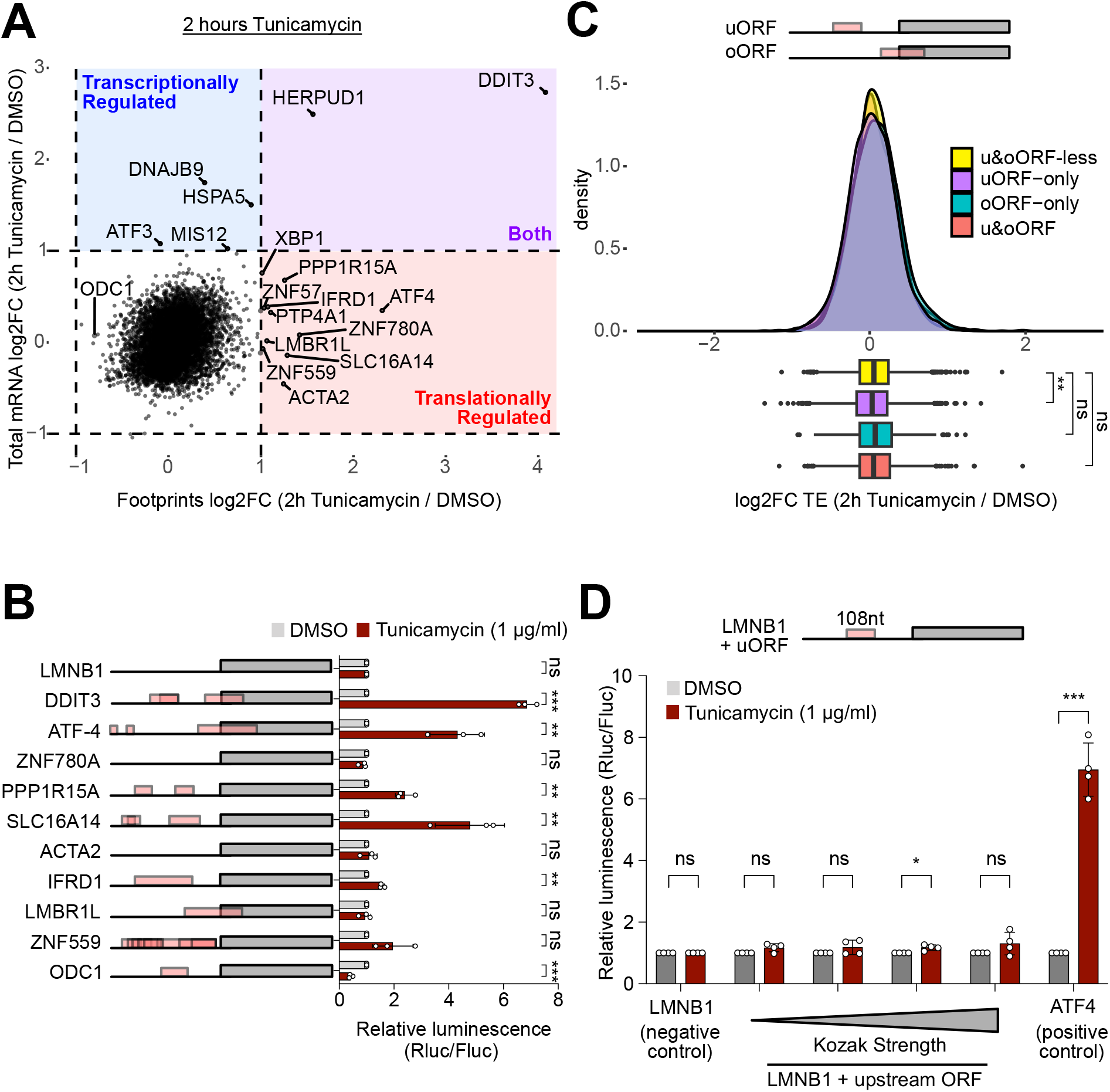
Ribosome footprinting identifies transcripts that are translationally induced upon induction of the Integrated Stress Response. (A) Ribosome profiling on HeLa cells treated with 1 µg/ml tunicamycin for 2-hours identifies only a handful of affected transcripts. Scatter plot showing change in translation (x-axis) versus change in mRNA levels (y-axis). (B) Only reporters bearing 5’UTRs with either uORFs or oORFs showed a translational response to tunicamycin treatment. n = 3 biological replicates. Error bars: standard deviation. Significance by unpaired, two-sided, t-test and adjusted for multiple testing. p-values: * p<0.05, ** p<0.01, *** p < 0.001, ns = not significant. (C) The simple presence of uORFs in an mRNA is a poor predictor of how translation of the mRNA will respond to ISR activation. Histogram of log2(fold change) of translation efficiency upon treatment of cells with tunicamycin (2h, 1 µg/ml) for different transcript groups. Box plots show median with lower and upper quartile. Whiskers: full data range. Significance by Kruskal-Wallis test followed by Dunn’s multiple comparison test. ns = not significant,* p<0.05, *** p<0.001. (D) Reporters containing synthetic uORF with Kozak sequences of different strengths are not induced in response to ISR activation. n = 3, biological replicates. Error bars: standard deviation. Significance by unpaired, two-sided, t-test and adjusted for multiple testing. p-values: * p<0.05, ** p<0.01, *** p < 0.001, ns = not significant.

### uORFs are necessary components of ISR-dependent translational activation, yet their presence alone is insufficient to predict translational upregulation

Previous studies described the presence of uORFs as a prerequisite for translational activation in response to the ISR (Andreev *et al.*, 2015; Sidrauski *et al.*, 2015). Indeed, in agreement with this, when we cloned luciferase reporters containing the 5′ UTRs of the top translationally activated transcripts, we found that only 5′ UTRs harbouring uORFs or oORFs were able to recapitulate the translational upregulation observed upon tunicamycin treatment (Fig. 1B). While translational activation of the other mRNAs likely relies on other features of the transcripts not present within their 5’UTRs (e.g. CDS or 3’UTR) this agrees with the observation that uORFs/oORFs are important for stress-induced translational activation. This also aligns with a previous report by Andreev et al. showing that almost all stress-resistant transcripts possess at least one efficiently translated uORF and that mutation of the uORFs of PPP1R15B and IFRD1 reduces their resistance to stress (Andreev *et al.*, 2015). To test this further, we selected two extreme examples from our dataset: ATF4, one of the most strongly translationally activated transcripts, and ODC1, one of the most strongly repressed. Mutation of all uORF/oORF start codons (AUG to UAC) within the ATF4 or ODC1 5′ UTRs abolished the up- or down-regulation, respectively (Suppl. Fig. 2D). Together, these results illustrate that uORFs can exert both activating and repressive effects on translation during the ISR.

To analyze whether all transcripts with uORFs are translationally regulated by tunicamycin, we classified the 7,685 transcripts we could detect into four categories: those containing only uORFs (2,495 transcripts, Kozak strength >= 50), those containing only oORFs (509 transcripts, Kozak strength >= 50), those containing both (1,161 transcripts), and those containing neither (3,520 transcripts). For these analyses, we took Kozak strength values from (Noderer *et al*, 2014) where they were termed TIS efficiency, and we retained only the transcript isoform with the highest translation efficiency for each gene. Comparison of translation efficiency values across these groups revealed no clear, global association between the presence or absence of uORFs/oORFs and the translational response to tunicamycin (Fig. 1C). This indicates that the vast majority of transcripts containing uORFs or oORFs does not become induced upon activation of the ISR. Hence the handful of transcripts that we found to be translationally induced by tunicamycin (Fig 1A) must have some special features in addition to simply having uORFs or oORFs. To test this experimentally, we introduced synthetic uORFs into a reporter carrying the LaminB1 5′ UTR, which lacks uORFs. Despite using Kozak sequences of different strengths on the uORF AUG, none of these uORFs imparted substantial induction in response to tunicamycin (Fig. 1D). Importantly, this excludes the possibility that ISR activation broadly increases leaky scanning of all uORFs. We therefore conclude that the mere presence of uORFs or oORFs is a poor predictor of translational activation and that the uORFs/oORFs that do cause activation in response to stress must have some special features. We therefore aimed to uncover what feature causes this induction upon activation of the ISR.

### Delayed initiator tRNA recruitment cannot explain translation induction of most transcripts upon stress

The original mechanistic explanation of how ATF4 translation increases during stress revolves around delayed recruitment of initiator tRNA. According to this model (Suppl. Fig. 3A), the ribosome translates the two ATF4 uORFs, and then continues scanning to reinitiate further downstream. Under non-stressed conditions, the scanning ribosome rapidly recruits a new ternary complex including the initiator tRNA. This causes it to initiate translation on the oORF, thereby bypassing the start codon of the ATF4 main ORF. Instead, when the ISR is activated, ternary complex levels are reduced, delaying reinitiation and allowing the ribosome to bypass the oORF AUG, initiating instead on the main ORF. Of note, this mechanism has been called into question (Zhou *et al.*, 2018) and recent work has shown that additional mechanisms involving ribosome queuing are at play (Smirnova *et al.*, 2024). We noticed that this mechanism involving delayed initiator tRNA recruitment cannot explain induction of the various transcripts we identified here. As previously described (Andreev *et al.*, 2015; Young *et al*, 2016), *IFRD1* and *PPP1R15A* have uORFs but no oORF (Fig. 1B). We identified here *SLC16A14* as an induced transcript (Fig. 1B), and it also has uORFs but no oORF. One transcript we found to be induced which has both uORFs and oORFs is *ZBTB45* (Suppl. Fig. 3B). Mutagenesis of this reporter revealed, however, that versions with only the uORFs but no oORF, or the other way around - only an oORF and no uORFs - are still inducible (Suppl. Fig. 3B). In sum, in most cases it is sufficient for a transcript to have only a uORF or an oORF to be inducible upon ISR activation, indicating that the induction does not occur due to delayed initiator tRNA recruitment.

### oORFs with the same degree of basal suppression show distinct translational responses upon ISR

To dissect the mechanism by which translation is induced upon tunicamycin treatment, we decided to study 5’UTRs containing oORFs but no uORFs. The reason is that uORFs can be overcome by either leaky scanning or reinitiation. In contrast, in the case of oORFs, the only way the main ORF can be translated is via leaky scanning, reducing the number of possible mechanistic explanations.

Both uORFs and oORFs with poor initiation contexts are unlikely to have regulatory functions because scanning ribosomes will probably ignore them altogether. Hence one plausible hypothesis was that transcripts containing uORFs/oORFs with strong initiation contexts, which cause stronger basal repression of the mORF under non-stressed conditions, might cause activation upon ISR induction. Although our synthetic reporters containing uORFs with strong Kozak contexts did not support this hypothesis (Fig. 1D), we could not exclude this was due to the synthetic nature of these reporters. We therefore cloned a panel of luciferase reporters containing endogenous 5’UTRs of transcripts with a single oORF per 5′ UTR (Suppl. Fig. 4A). Measurement of the activity of these reporters in non-stressed conditions (Fig. 2A) revealed differing levels of repression of luciferase translation, as expected by the fact that the oORF AUGs have initiation contexts of different strengths. Also as expected, oORFs that were poorly translated, and hence caused little repression of luciferase translation under non-stressed conditions (e.g. JOSD2 or FARP2) also caused no translational induction in response to stress (Fig. 2B). One of the reporters, carrying the 5’UTR of DCP2, showed strong induction upon tunicamycin treatment (Fig. 2B). The oORF of DCP2 shows a strong inhibitory effect on basal luciferase activity (Fig. 2A), indicating that the oORF is well translated and traps the majority of initiating ribosomes, in agreement with our hypothesis. DCP2 activation was specific to the ISR, as treatment with the mTOR inhibitor Torin failed to induce reporter activation (Suppl. Fig. 4B). Unexpectedly, however, other reporters with equally well translated oORFs (e.g. RNFT1, Fig. 2A) did not show induction in response to tunicamycin (Fig. 2B). We conclude that the start codon initiation context is not sufficient to predict translational activation during the ISR.

**Figure 2.**
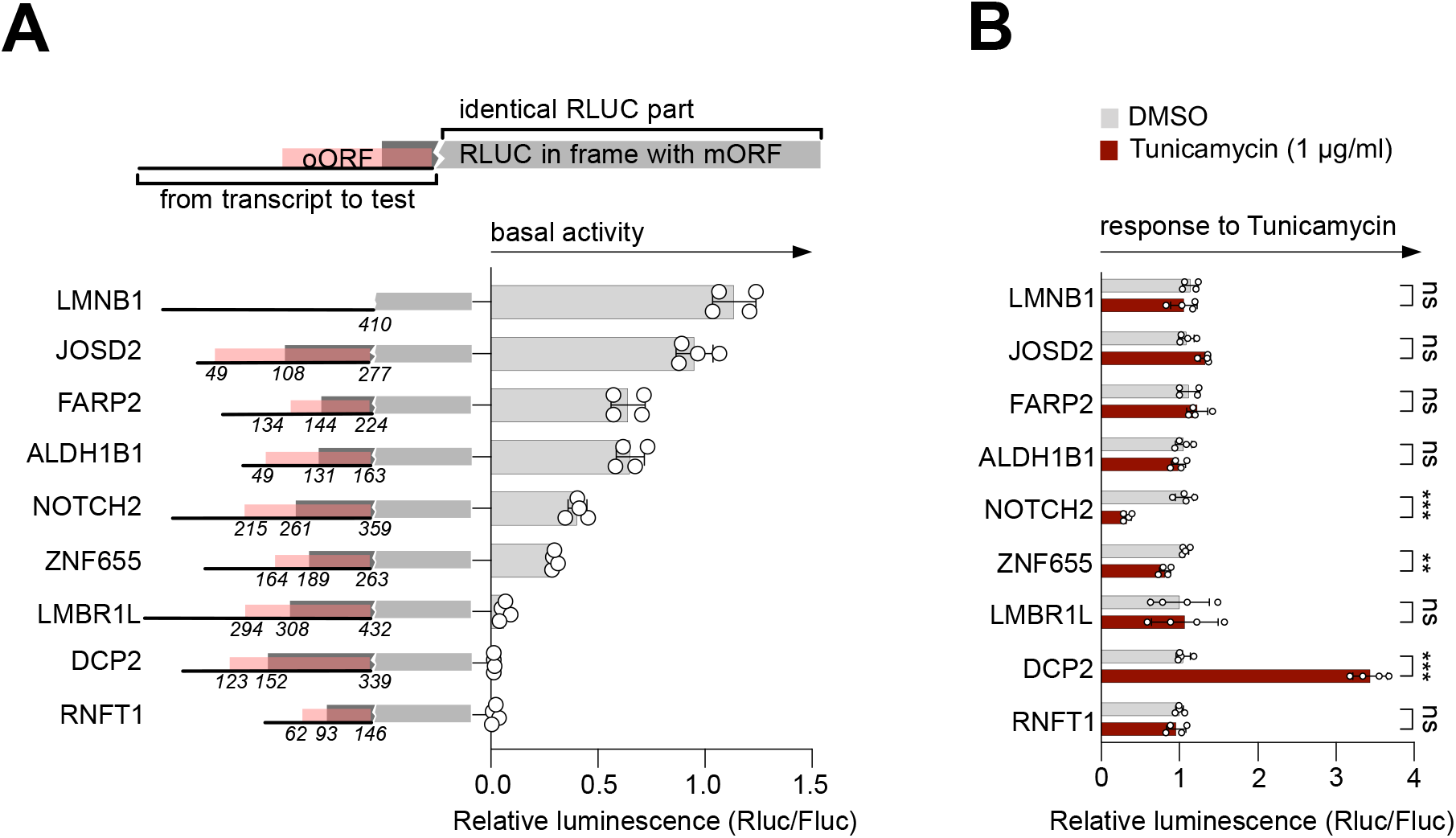
A reporter bearing the 5’UTR of DCP2 is strongly activation upon ISR activation. (A) Luciferase activity in basal, non-stressed conditions, of reporters bearing 5’UTRs of different genes. Schematic representation of cloned reporters, with position of mORF and oORF start, as well as oORF stop is indicated to the left. n = 4, biological replicates. (B) A reporter bearing the DCP2 5’UTR is strongly induced upon induction of the ISR. Reporter response to tunicamycin treatment (16h) for the same reporters as in (A). Significance by unpaired, two-sided, t-test and adjusted for multiple testing. Error bars: standard deviation. p-values: * p<0.05, ** p<0.01, *** p < 0.001, ns = not significant.

### Kozak’s strength is not altered between physiological and stress conditions

A possible mechanistic hypothesis to explain how stress could lead to activation of mORF translation is if stress would more potently inhibit initiation on the oORF compared to the mORF due to their different Kozak sequences. For instance, if an oORF were to have a Kozak sequence whose strength drops strongly upon stress, whereas the mORF has a Kozak sequence that only drops mildly, this would lead to a relative increase of translation initiation on the mORF compared to the oORF (Suppl. Fig. 5A). To test whether stress differentially inhibits initiation from start codons with different Kozak sequences, we performed a Kozak library screen. Using oligo cloning, we generated a panel of Renilla luciferase (RLuc) reporters containing a synthetic oORF with Kozak sequences of varying strengths inserted into the LaminB1 5′ UTR (Suppl. Fig. 5B). Importantly, each plasmid also encoded a Firefly luciferase reporter driven by an oORF-less LaminB1 5′ UTR, enabling internal normalisation. All transfections were normalised to a control in which both Renilla and Firefly luciferase were guided by the oORF-less LaminB1 5′ UTRs. In total, we cloned 124 Kozak variants. A previous study measured Kozak sequence strength using a reporter-based approach combined with FACS sorting under non-stressed conditions (Noderer *et al*, 2014). Our measurements closely recapitulated these findings, with stronger Kozak sequences causing stronger suppression of translation on the RLuc mORF (Suppl. Fig 5C). Treatment with tunicamycin, however, did not alter the relative strength of different Kozak sequences. Suppl. Fig 5D shows a scatter plot where each dot represents one reporter, with its expression level in non-stressed conditions on the x-axis and its expression level upon tunicamycin treatment on the y-axis. If some Kozak sequences would be differentially affected by stress compared to the others, they would lie off the diagonal, which was not the case (Suppl. Fig. 5D). Therefore, we conclude that stress equally represses translation initiation on all AUGs regardless of their Kozak sequence.

### Induction of DCP2 requires its oORF

To perform a detailed dissection of the underlying mechanism of induction, we selected to study in depth the DCP2 reporter, which has one oORF and is robustly induced upon tunicamycin treatment. We first confirmed that mutation of the oORF start codon (AUG to UAC) completely abolished ISR-dependent induction, demonstrating that translation of the oORF is required for this effect (ΔoORF, Fig. 3A). We then proceeded to systematically test the features of the DCP2 5’UTR from the 5’ end to the 3’ end by removing or altering them one by one.

**Figure 3.**
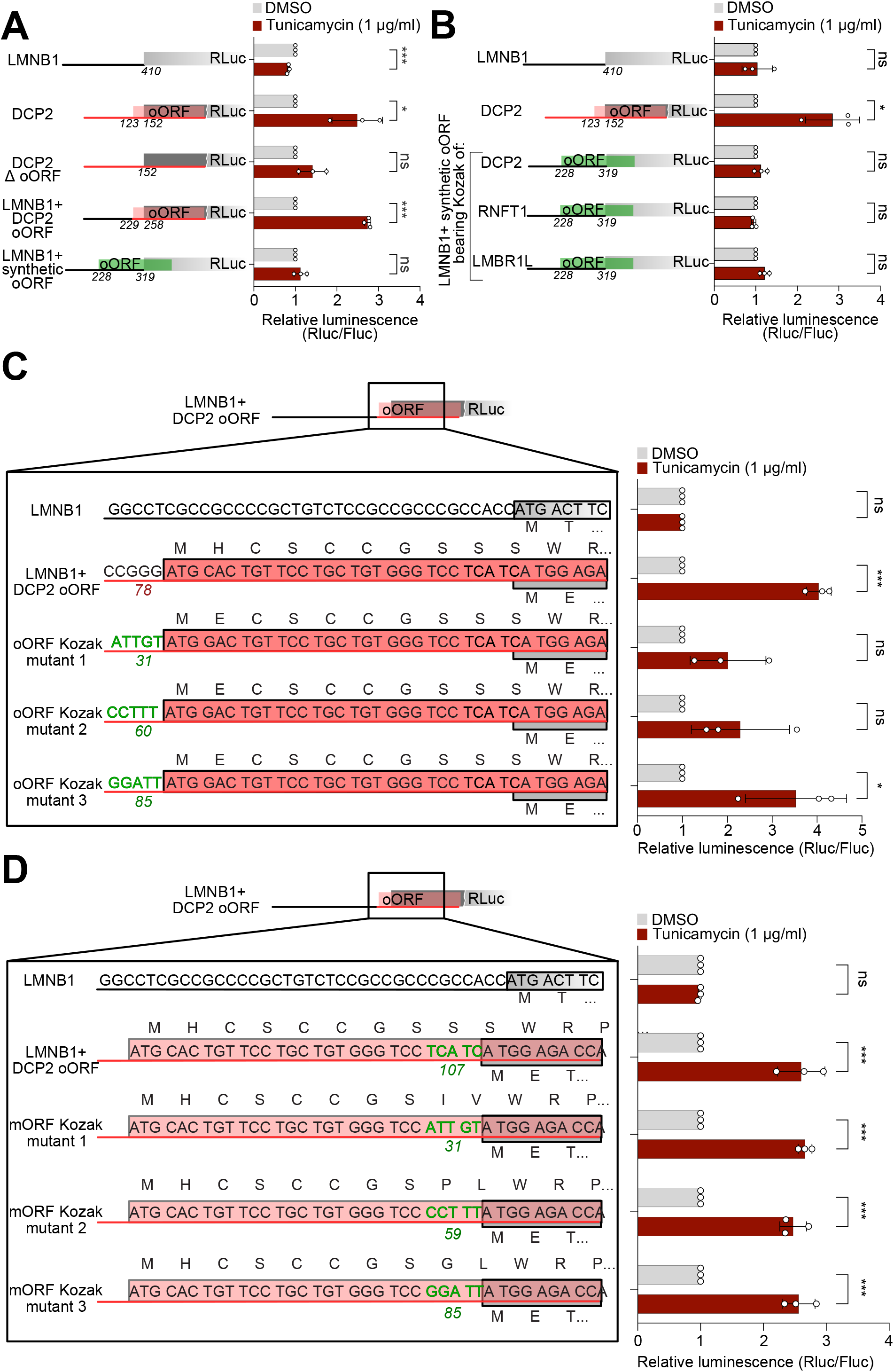
The DCP2 reporter requires a strong Kozak on the oORF for inducibility. (A) Induction of DCP2 relies on presence of an oORF. Mutation of the oORF start codon to UAC abolishes induction, while substitution of the entire sequence upstream of the oORF with the LMNB1 leader does not affect inducibility. (B) The Kozak sequence of the DCP2 oORF alone is insufficient to confer inducibility. Luciferase reporters bearing a synthetic oORF with the Kozak sequence of either DCP2, RNFT1 or LMBR1L are all not induced upon activation of the ISR. (C) Robust induction of the DCP2 reporter requires a strong Kozak context on the oORF start codon. The Kozak sequence of the DCP2 oORF was mutated to sequences with varying strengths (indicated numerically with arbitrary units on the left, taken from (Noderer *et al.*, 2014)) and transfected into HeLa cells treated with either DMSO or tunicamycin. Strong induction was observed only for reporters containing a strong Kozak sequence. (D) The strength of the Kozak sequence on the main ORF of the reporter has little impact on DCP2 inducibility. Mutation of the main ORF Kozak sequence to sequences of various strengths (indicated on the left) doesn’t affect reporter inducibility. Kozak strength are taken from (Noderer *et al.*, 2014). All panels: n=3 Significance by unpaired, two-sided, t-test and adjusted for multiple testing. Error bars: standard deviation. p-values: * p<0.05, ** p<0.01, *** p < 0.001, ns = not significant.

To assess whether any feature upstream of the DCP2 oORF is required, we generated a chimeric reporter in which the region upstream of the DCP2 oORF Kozak sequence was replaced with the corresponding region from the negative control LaminB1 5′ UTR. Notably, this chimeric construct retained an induction level comparable to that of the full-length DCP2 5′ UTR reporter (Fig. 3A) indicating that sequences upstream of the oORF are not required for inducibility. In agreement with this, the converse experiment where we retained the sequence upstream of the DCP2 oORF but swapped the oORF for a synthetic oORF abolished induction (Fig. 3A) indicating that there is something special about the oORF sequence which imparts stress inducibility.

We next tested whether the oORF Kozak sequence was sufficient to confer stress-induced activation. To this end, we introduced the DCP2 oORF Kozak sequence into a synthetic oORF reporter. As controls, we also generated synthetic oORF reporters containing Kozak sequences derived from the RNFT1 and LMBR1L oORFs, which did not show translational upregulation in our earlier assays (Fig. 3B). However, the DCP2 oORF Kozak was insufficient to induce reporter activation upon tunicamycin treatment (Fig. 3B). In agreement with this, replacement of the oORF Kozak sequence with another Kozak sequence permitted the reporter to retain inducibility. We mutated the DCP2 oORF Kozak context to sequences of varying strengths, and observed that even a weak Kozak context permitted stress-induced activation of the DCP2 reporter, although maximal induction required a strong Kozak context (Fig. 3C, Suppl. Fig 6B). Consistent with this observation, analysis of Kozak strengths associated with u/oORFs causing translational activation - identified either by us or by others (Andreev *et al.*, 2015; Young *et al.*, 2016) - revealed that regulatory, suppressive uORFs frequently harbour strong Kozak sequences (Suppl. Fig. 6A). Thus, for optimal activation the Kozak sequence of the oORF should be strong, but there is nothing special about the exact sequence of the DCP2 oORF because it can be exchanged with another strong Kozak sequence.

We next analysed the sequence of the DCP2 oORF and found that secondary-structure prediction indicated the presence of a hairpin (Suppl. Fig. 6C). To test whether this hairpin contributes to inducibility, we introduced mutations that disrupt pairing in the stem while maintaining the amino acid coding of the oORF (Δ, Suppl. Fig. 6C). Since this mutant still retained some pairing in the stem, we also generated a more drastic mutant that completely disrupted the hairpin and also changed the oORF amino acid coding (ΔΔSuppl. Fig. 6C). Neither mutant, however, exhibited reduced inducibility compared to the wild-type reporter (Suppl. Fig. 6D), leading us to conclude that the predicted secondary structure of the DCP2 oORF does not have a major impact on its stress-induced translational activation. Furthermore, since the ΔΔ mutation also changed part of the amino acid sequence of the oORF between the oORF start codon and the mORF start codon, it suggests that this region of the oORF upstream of the mORF start codon may not be critical for inducibility.

We next examined whether the Kozak sequence of the DCP2 main ORF contributes to reporter inducibility. Substitution of the native mORF Kozak sequence with sequences of varying strengths only had a minor effect on DCP2 induction upon tunicamycin treatment, indicating that mORF Kozak identity is not a major determinant of this response (Fig. 3D).

In summary, DCP2 inducibility requires the oORF but does not appear to require any specific sequence of the oORF upstream of the mORF start codon.

### The 3’ end of the DCP2 oORF is required for ISR-mediated induction

Given that stress inducibility for another tunicamycin-induced transcript, DDIT3 (also known as CHOP), was shown to depend on the identity of the peptide encoded by the uORF (Young *et al.*, 2016; Young & Wek, 2016), we next asked whether the coding sequence of the DCP2 oORF plays a functional role. We first tested the oORF region that is upstream of the mORF start codon, by replacing it with the equivalent regions from the synthetic oORF or the RNFT1 oORF, both of which are not stress-inducible (Fig. 2A, 3A). These chimeric reporters (Suppl. Fig. 7A) were still stress-inducible (Fig. 4A-B), indicating that the sequence of the oORF upstream of the mORF AUG is not critical for inducibility.

**Figure 4.**
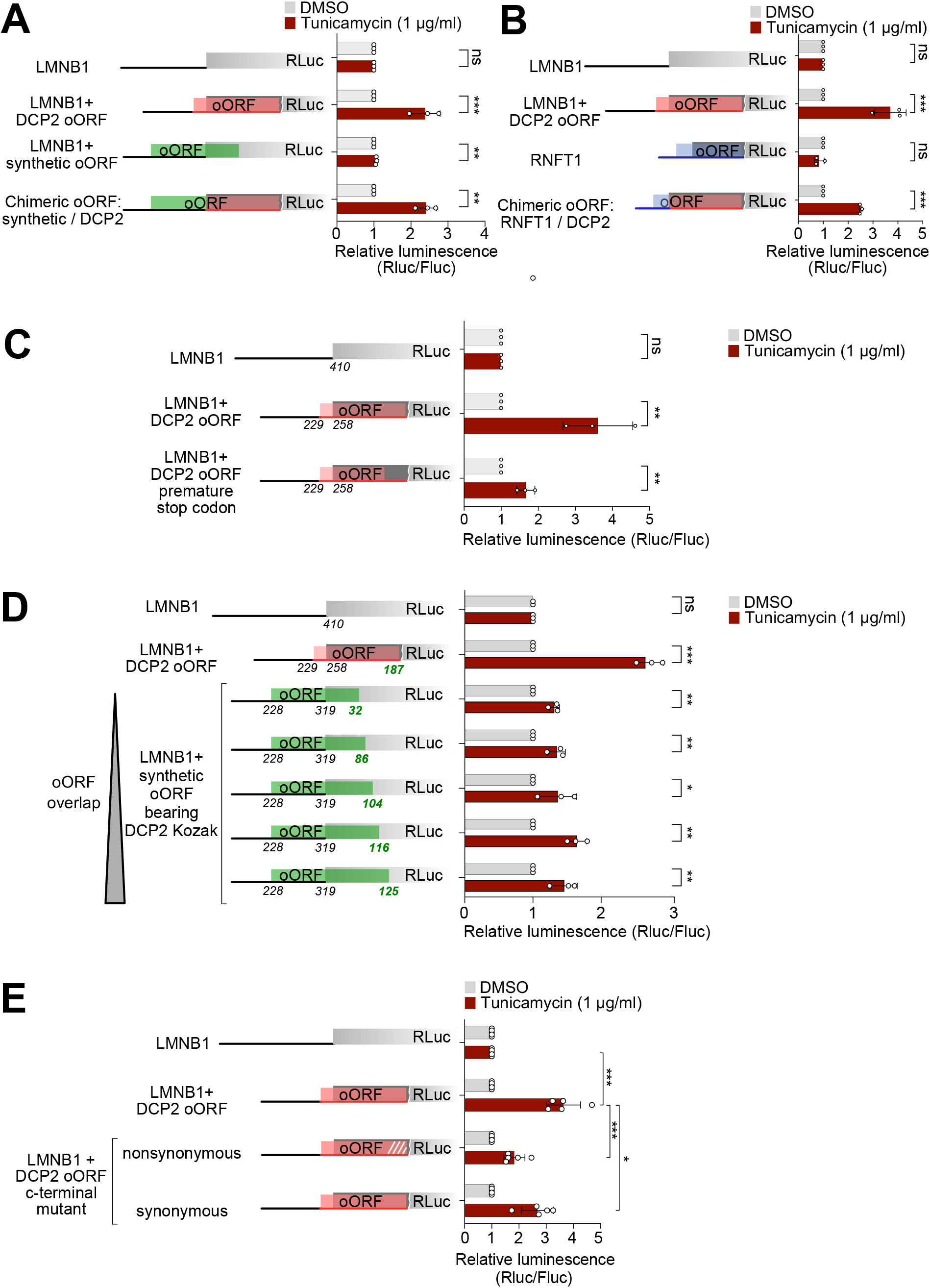
DCP2 induction depends on an element at the 3’end of the oORF. (**A-B**) The region of the DCP2 oORF upstream of the main ORF is dispensable for ISR-mediated induction. Chimeric constructs, in which the region of the DCP2 oORF that is upstream of the main ORF was swapped with the equivalent portion of either a synthetic oORF (A) or the RNFT1 oORF (B), exhibited induction levels comparable to the full-length DCP2 reporter. (C) Shortening of the DCP2 oORF by introducing a premature termination codon blunts inducibility of the reporter. The premature stop codon in the oORF was introduced without affecting the protein coding of the main ORF, as indicated in Suppl. Fig. 7B (D) A large overlap between the oORF and the main ORF is not sufficient to impart inducibility. Synonymous mutations were introduced into the RLuc CDS to extend the overlap of a synthetic oORF, yet none of the reporters displays the same inducibility as the DCP2 reporter. (E) Changes in the codons at the 3’ end of the DCP2 oORF reduce reporter inducibility. The precise mutations are indicated in Suppl. Fig 7E. Panels A-D: n=3 Significance by unpaired, two-sided, t-test and adjusted for multiple testing. Error bars: standard deviation. p-values: * p<0.05, ** p<0.01, *** p < 0.001, ns = not significant. Panel D: n=5 Significance by two-way ANOVA with Dunnett’s multiple comparison test. Error bars: standard deviation. p-values: * p<0.05, *** p < 0.001.

We next turned our attention to the part of the oORF that is downstream of the mORF start codon. We introduced a premature stop codon within the DCP2 oORF without altering the amino acid coding of the mORF (Suppl. Fig. 7B). This mutation strongly blunted the induction of the DCP2 reporter upon tunicamycin treatment (Fig. 4C). Notably, introduction of this stop codon also derepressed basal reporter expression to a level comparable to that observed upon mutation of the oORF start codon (Suppl. Fig. 7C), indicating that the 3’ end of the oORF is repressive for mORF translation. This experiment does not distinguish whether the relevant regulatory feature is the C-terminal sequence of the oORF peptide, or the length of overlap between the oORF and the mORF, which is particularly long in DCP2. To test whether the key feature is the length of overlap, we generated a series of reporters with a synthetic oORF with increasing overlap into the Renilla luciferase mORF, however, this did not result in substantial stress-induction (Fig. 4D).

We then analyzed the oORF sequence downstream of the mORF AUG. Interestingly, the amino acid sequence encoded by the DCP2 oORF is highly conserved across species, suggesting evolutionary constraint and a potential regulatory function (Suppl. Fig. 7D). Evolutionary conservation of the overlapping mORF amino acid sequence may contribute to this conservation, however variance in the wobble positions of the mORF would be expected to disrupt conservation of the oORF at the amino acid level. To directly test the importance of the oORF sequence, we exploited wobble-base substitutions at the third codon position to alter the amino acid sequence of the C-terminal region of the DCP2 oORF without affecting the amino acid sequence of the mORF (Suppl. Fig. 7E). These mutations resulted in a marked blunting of the stress-induction (“nonsynonymous”, Fig. 4E).

### The 3’end of the DCP2 oORF contains an inhibitory element

We envisioned two possible molecular mechanisms to explain the inducibility of DCP2. A ’leaky scanning’ competition model would propose that ribosomes initiate translation either on the oORF or on the mORF. Hence upon ISR activation, initiation on the mORF would increase because initiation on the oORF decreases, and therefore the 3’ end of the DCP2 oORF somehow reduces initiation on itself (Suppl. Fig. 8A). An alternative model proposes that the 3’end of the oORF contains an inhibitory element which blocks translation of both the oORF itself and the mORF, and this inhibition is relieved in response to the ISR. According to this model, translation of both the oORF and the mORF should increase upon ISR activation (Suppl. Fig. 8A). Hence the two models have diametrically opposed predictions what should happen to oORF translation in response to stress. To test this, we added an HA-tag to the 5’ end of the DCP2 oORF and assayed what happens to levels of the HA-oORF peptide in response to tunicamycin (Suppl. Fig. 8B). This revealed that activation of the ISR increases translation of the HA-oORF peptide, rather than decreasing it, consistent with the model postulating an inhibitory element. Consistent with this, fusing the DCP2 oORF in frame with RLuc strongly reduced RLuc levels (Suppl. Fig. 8C), again indicating that the DCP2 oORF contains an inhibitory element. Finally, this in-frame fusion increased in expression upon treatment with tunicamycin (Suppl. Fig. 8D). Altogether, these data indicate that in response to ISR activation, expression of both the oORF and the mORF of DCP2 increase in parallel, indicating that there is a repressive element at the 3’ end of the DCP oORF which is relieved upon stress induction.

Interestingly, in a translatome-wide screen (Han *et al*, 2026) this 3’ end of the DCP2 oORF was identified as a strong ribosome pausing site, ranked at position 52 genome-wide when sorted by pausing strength. This suggests that under non-stressed conditions the DCP2 oORF causes ribosomes to stall. Since this position is located within the mORF, this also inhibits translation of the main ORF. When cells experience stress, either the stalling is reduced or global translation rates drop so that the stalling becomes proportionally less relevant, allowing the main ORF to be translated (Fig. 5A). If this is the case, then this mechanism should also work if the DCP2 oORF is shifted entirely into the 5’UTR as a uORF, because stalling on a uORF should also block translation of the downstream mORF. To test this, we generated a synthetic reporter in which the DCP2 oORF was inserted into the LaminB1 5′ UTR as a uORF (Suppl. Fig. 9A). (To prevent the formation of an alternative reading frame derived from the DCP2 mORF start codon that is present inside the oORF, this start codon was mutated to UAC, thereby ensuring that the resulting uORF was translated only in one frame.) As predicted, this chimeric reporter showed induction upon tunicamycin treatment (Suppl. Fig. 9A). Hence the regulatory activity of the DCP2 oORF also functions if it is present as a uORF, fitting with the hypothesis that it acts as a stalling site.

**Figure 5.**
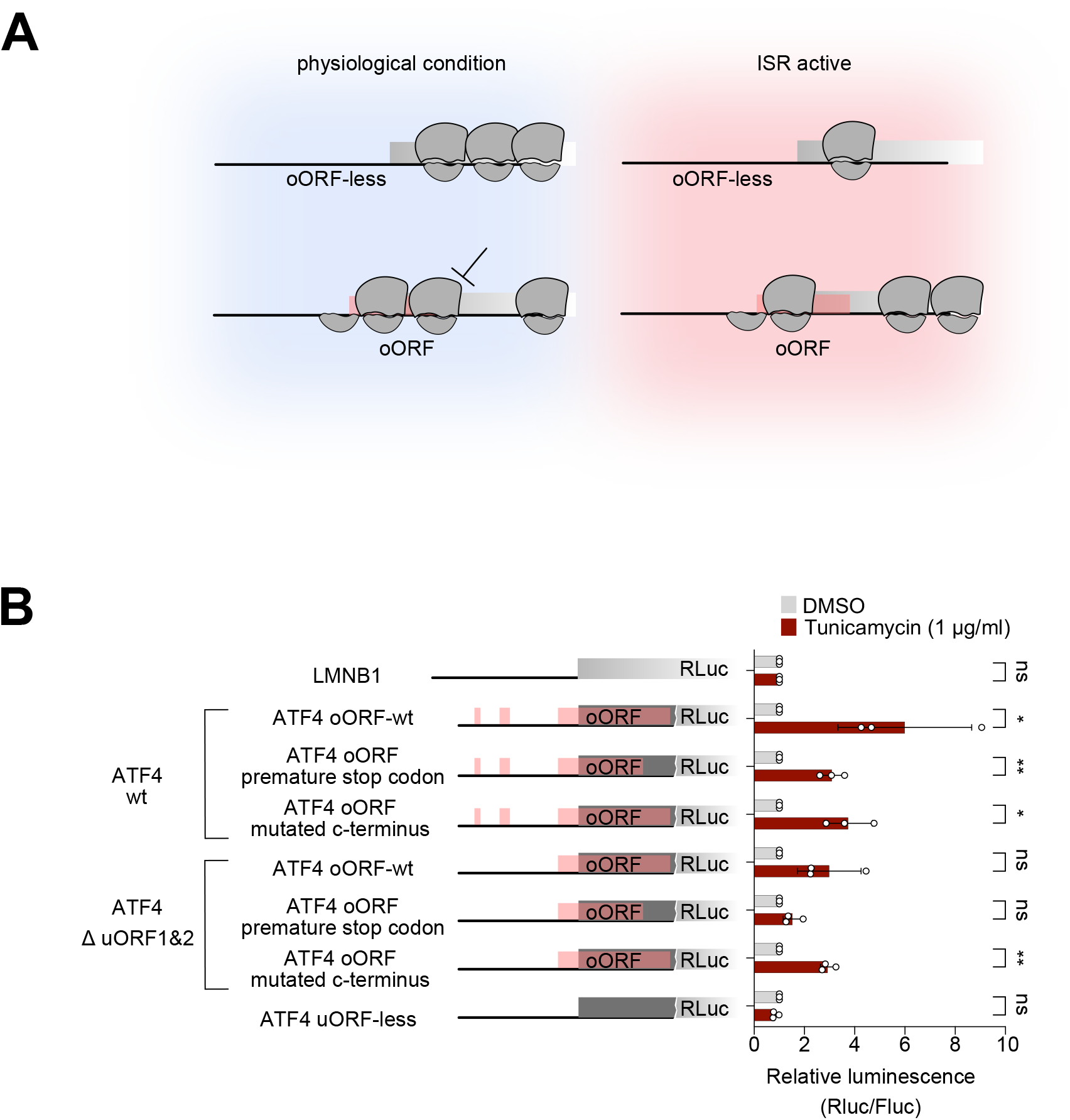
Translational induction in response to ISR activation may function via relief of a repressive mechanism. (A) Schematic representation of the proposed model. Under basal conditions, ribosomes mainly translate the oORF, which has a repressive element at its 3’ end, leading to ribosome stalling and repression of translation of both the oORF and the main ORF. When the integrated stress response is activated, the repressive activity of this element is suppressed, leading to increased translation of both the oORF and the mORF. (B) Induction of an ATF4 reporter is possible even when the uORFs are mutated, leaving only the overlapping ORF. Induction of the reporter is blunted when a premature stop codon in the oORF is introduced, or the amino acid sequence of encoded by oORF peptide is altered. n=3 Significance by unpaired, two-sided, t-test and adjusted for multiple testing. Error bars: standard deviation. p-values: * p<0.05, ** p<0.01, *** p < 0.001, ns = not significant.

The stalling may be caused either by suboptimal codons which are slow at being translated, or by an interaction of the nascent polypeptide with the ribosomal exit tunnel. To distinguish these two options, we introduced synonymous mutations in the wobble positions of the oORF, which change the oORF codons but not the encoded polypeptide (Suppl. Fig. 7E). This caused an intermediate drop in the induction of the reporter in response to tunicamycin (Fig. 4E), suggesting that both the identity of the codons and the encoded protein contribute to inducibility. Future work will be required to study this in more depth.

### ATF4 induction involves mechanisms in addition to delayed recruitment of initiator tRNA

Finally, we asked if we could extrapolate some of our findings for the DCP2 reporter to other stress-induced genes. We decided to focus on the classical ISR target ATF- 4. To address this, we generated a series of ATF4 reporter constructs in which we mutated the AUG start codons of the two uORFs to UAC and/or mutated the oORF. Consistent with the DCP2 results, we found that the reporter containing only the ATF4 oORF remains inducible upon tunicamycin treatment (Fig 5B, Suppl. Fig. 9B). This is consistent with data in the original paper identifying ATF4 as a stress-induced transcript (Vattem & Wek, 2004), where Fig. 4A shows that the reporter lacking the uORFs is still induced. Our results are also consistent with a more recent, in-depth dissection of the ATF4 induction mechanism (Smirnova *et al.*, 2024). Since no re-recruitment of initiator tRNA is necessary in this configuration when the uORFs are gone, the remaining inducibility suggests additional, oORF-intrinsic mechanisms are at play besides delayed re-initiation. Indeed, shortening the oORF or mutating its C- terminal amino acid sequence (Suppl. Fig. 9C) reduced inducibility, both in the presence and absence of uORFs (Fig. 5B), indicating that oORF length and peptide/codon sequence contribute to stress-induction. In sum, there are close parallels between the DCP2 oORF and the ATF4 oORF in that the 3’ends of both oORFs contribute to inducibility. In the case of the ATF4, the element is a stem-loop (Smirnova *et al.*, 2024) whereas in the case of DCP2 it is a ribosome stalling site.

## Discussion

We identify here DCP2 as a novel stress-regulated transcript that is translationally induced in response to activation of the Integrated Stress Response (ISR). Since DCP2 encodes for an mRNA de-capping enzyme, it may be involved in de-capping mRNAs that are not needed upon stress. Increased mRNA turnover would allow replacement of the mRNAs that were transcribed prior to stress with the newly transcribed mRNAs. We find that induction of DCP2 is dependent on a sequence encoded by the 3’end of the DCP2 overlapping ORF (oORF), which corresponds to a strong ribosome collision site (Han *et al.*, 2026).

We propose that inhibitory elements in the 3’ end of uORFs or oORFs might be a general mechanism for stress-inducibility. In the case of DCP2, it corresponds to a stalling site in the oORF. For ATF4, the oORF contains a stem-loop that is inhibitory (Smirnova *et al.*, 2024). In the case of CHOP/DDIT3 it is a ribosome stalling site in the oORF (Young *et al.*, 2016), analogous to DCP2. Finally, for GADD34/PPP1R15A, the mRNA contains a uORF with an inhibitory Pro-Pro-Gly sequence (Young *et al*, 2015).

Whether the inhibitory element is in a uORF on an oORF appears not to be very important - we found that the DCP2 inhibitory element from the oORF can be transferred into a uORF and it still imparts inducibility. If this is the case, one open unresolved question is how stress inhibits or dampens this inhibitory activity? One possible mechanism could be if the inhibitory element consists of poorly-translated codons. Activation of the ISR would lead to a global repression of translation, thereby increasing the availability of rare tRNAs, allowing the inhibitory element to be translated more efficiently. Future work in this direction might uncover interesting molecular mechanisms.

## Material and Methods

### Cell lines, culture conditions, and treatments

All experiments were performed in a HeLa cell line that was authenticated using SNP typing. Cells were cultured in DMEM (Gibco 41965039) supplemented with 10% fetal bovine serum (FBS) (Sigma, S0615) and 100 U/ml Penicillin/Streptomycin (Gibco 15140122). For the indicated experiments, cells were treated either with DMSO, or with 1 µg/ml tunicamycin (PanReac AppliChem, A2242,0005), ISRIB (SML0843-25M) or 250nM Torin (Sigma, 475991-10MG) for the indicated periods of time.

### Cloning

Firefly (pAT1620) and Renilla luciferase (pAT1618) constructs under the control of the LMNB1 5′UTR were described previously (Schleich *et al*, 2017). The 5′UTRs of transcripts to be tested were PCR-amplified from HeLa cDNA using the oligonucleotides listed in Table S2. For reporters containing overlapping ORFs, the native initiation context of both the overlapping and main ORFs was preserved. The reverse primer was designed to fuse the endogenous main ORF in-frame with the Renilla luciferase ORF. PCR products were gel-purified, digested with HindIII and Bsp119I, and cloned into pAT1618 using the same restriction sites. The cloned 5′UTR variants are listed in Table S3 along with their corresponding transcript IDs. Reporters containing uORFs with varying Kozak sequence strengths were generated as previously described (Bohlen *et al.*, 2023). uORF variants, secondary structure mutants, Kozak sequence mutants, and chimeric constructs were produced by site-directed mutagenesis PCR using oligonucleotides listed in Table S2. The resulting PCR products were cloned back into the backbone between HindIII and Bsp119I sites. For subcloning the DCP2 oORF into the LMNB1 5′UTR, an LMNB1 reporter containing Kpn2I and BshT1 sites (Schleich *et al*, 2014) was used. The DCP2 region containing the oORF was amplified using primers incorporating Kpn2I and BshTI restriction sites and cloned into the corresponding sites of the LMNB1 reporter. Following this step, the start codon of the main ORF (mORF) was mutated using site-directed mutagenesis. All constructs were validated by Sanger sequencing and are available at the European Plasmid Repository.

### Ribosome profiling

HeLa cells were seeded at 2 million cells per 15 cm dish in 20 ml of growth medium one day before harvesting. The following day, cells were treated either with DMSO or 1 µg/ml tunicamycin for 2 hours. After treatment, cells were quickly washed with ice-cold 1xPBS supplemented with 10 mM MgCl2 and 800 µM Cycloheximide. After the wash, all residual solution was removed by gentle taping of the 15cm dish on its side, followed by cell lysis with 150 µl of the following buffer: 0,25 M HEPES pH 7.5, 50 mM MgCl2, 1 M KCl, 5% NP40, 1000 μM Cycloheximide. Cells were scraped into an Eppendorf tube and the lysate was clarified by centrifugation at 15.000xg for 10 minutes at 4°C. The concentration of lysate was estimated using a nanodrop spectrophotometer, measuring RNA content against a water-blanked control. 150 ul of lysate was used for the total RNA preparation with the RNeasy kit (Qiagen, cat. No. 74106). The remaining lysate was used for treatment with RNaseI (100 Unit per 120µg of lysate) on ice for 30 minutes. Following digestion, the lysates were loaded on a 15- 65% sucrose gradient, which was prepared in advance with the use of a Biocomp Gradient Master. The lysate was ultracentrifuged for 3 hours at 35000 rpm in a Beckman Ultracentrifuge with a SW40Ti rotor. To collect the 80S fraction the gradient was separated on a Biocomp Gradient Profiler system. The collected 80S fractions were used for RNA extraction with acid-phenol as described previously (Roiuk *et al.*, 2025). The integrity of extracted RNA was analysed on a Bioanalyser. The sample was then divided into two equal parts: one half underwent rRNA depletion using the Illumina Ribo-Zero Gold kit (now discontinued), while the other half was used directly for size selection. To size-select footprints, RNA extracted from the 80S peak was separated on a 15% Urea-Polyacrylamide gel and fragments of 25-35 nucleotides were gel-extracted. For this, the footprint-containing gel pieces were broken into small pieces with gel smasher tubes. 0.5 ml of 10 mM Tris pH 7 were added to the smashed gel pieces and the suspension was incubated at 70°C for 10 minutes with shaking. The mix was briefly centrifuged and the supernatant was used for RNA precipitation by isopropanol. Purified footprints were phosphorylated by means of T4 PNK (NEB) for 1 hours at 37°C in PNK buffer supplemented with 10 mM ATP. After this, the footprints were again precipitated and purified using isopropanol. To estimate the quality of footprints, RNA was run on an Agilent Bioanalyzer small RNA chip, followed by library preparation with the Next-Flex small RNA v.4 kit protocol (Perkin Elmer, NOVA-5132-06), in accordance to manufacture recommendations. Total RNA libraries were prepared using the Illumina TruSeq Stranded library preparation kit. The quality of libraries was checked on a Bioanalyser with the use of a High sensitivity DNA kit (Agilent, 5067-4626). The libraries were sequenced on an Illumina Next-Seq 550 system. The experiment was conducted in duplicate. However, given that each 80S sample was split into two fractions—with and without rRNA depletion— footprint libraries resulted in two DNA libraries per sample. All libraries were included in the data analysis.

### Data Analyses of Ribosome profiling

Reads were trimmed to remove adaptors and randomized nucleotides derived from use of the Nextflex kit with cutadapt software. By use of bowtie2 the reads aligning to tRNA or rRNA were removed. All remaining reads were mapped to the human transcriptome (Ensemble transcript assembly 94) and genome (hg38) using BBMap software, with multiple mapping allowed. The reads mapping to the coding sequences were quantified with lab-based software written in C (https://github.com/aurelioteleman/Teleman-Lab). For each transcript the value of reads per kilobase of coding sequence was calculated and only transcripts with values more or equal of 2.5 were used for the subsequent analysis. Metagene profiles were built with the custom-made software written in C. For the metagene profile of uORF stop codons, only transcripts with a space of more than 50 nucleotides between uORF and main ORF were used. To obtain values of translation efficiency, log2 fold changes, and adjusted p values the DESeq2 software package was used. Ribosome profiling data have been deposited at NCBI GEO with accession number GSE336812.

### Preparation of cell lysates with RIPA buffer

Cells were seeded at a density of 0.5 million cells per well of 6-well plate. Following possible treatment with drugs, cells were washed briefly with PBS, and then lysed with 120µl of RIPA buffer supplemented with 20U of Benzonase (Merk Millipore, 70746-3), protease (Sigma, 4693159001) and phosphatase (Sigma, 4906837001) inhibitors. Cells were collected by scraping followed by lysate clarification by centrifugation at 4°C for 10 minutes at 20.000g. The protein concentration was measured by Pierce BCA Protein assay kit (Life technologies, 23225). Protein concentration was equalised between samples and mixed with 5x Laemmli buffer (1/5 of the final volume). Samples were incubated at 95°C for 5 minutes and then loaded on an SDS-PAGE gel.

### Western blotting

Cell lysates were separated on SDS-PAGE gels, and transferred to a nitrocellulose membrane with 0.4 µm pore size (Amersham, 10600002) by wet transfer, and the membrane was blocked in 5% skim milk / PBST for 1 hour. Membranes were then incubated overnight in primary antibody solution (5% BSA / PBST) at 4°C. On the following day membranes were washed three times 15’ in PBST and incubated in secondary antibody (1:10000 in 5% skim milk / PBST) for 2 hours at room temperature, followed by three 15 minutes washes in PBST. Finally, chemiluminescence was detected with ECL reagents (Thermo Schientific, 32109) and imaged with a Biorad ChemiDoc imaging system. Antibodies used for immunoblotting are listed in Table S4.

### Dual-luciferase translation reporter assay

Cells were seeded in 96-well plates at a density of 8,000 cells per well one day prior to transfection. The following morning, cells were transfected using Lipofectamine 2000 with 100 ng of Renilla luciferase plasmid and 100 ng of Firefly luciferase plasmid per well. For the Kozak screen, cells were transfected with 200 ng of a single plasmid containing both Firefly and Renilla luciferase reporters driven by independent promoters. Six hours post-transfection, the medium was replaced with fresh medium supplemented with either DMSO or 1 µg/mL tunicamycin. 16 hours after the medium change, luciferase activity was measured using the Promega Dual-Luciferase Assay System (Promega, E1910) according to the manufacturer’s instructions.

## ACKNOWLEDGEMENTS

We thank the DKFZ Genomics Core Facility for next-generation DNA sequencing.

## CONFLICT OF INTEREST

Authors declare no competing interests.

**Supplementary Figure 1 Two hours of 1 µg/ml tunicamycin treatment is sufficient to induce the Integrated Stress Response and to blunt translation.** (**A-C**) Western blot assessment of ISR activation during a 1 µg/ml tunicamycin treatment time course of HeLa cells. An increase in the phosphorylation of eIF2α is detected already after 1 hour of treatment, with a substantial increase in ATF-4 levels after 2 hours. Ratio of p-eIF2α to total eIF2α is quantified in (B). The ratio of ATF4 levels to tubulin levels are quantified in (C). Error bars: standard deviation.

(D) Polysome profiles of HeLa cells treated with 1 µg/ml tunicamycin for different time points show substantial repression of translation at 2 hours of treatment. Lysates were separated on a sucrose gradient. The polysome/80S ratio is indicated.

(E) Detection of the spliced variant of Xbp1 coincides with the dynamics of eIF2a phosphorylation. Spliced variant at the zero time point was not detectable. Error bars: standard deviation. Significance by unpaired, two-sided t-test. *** p<0.001

(F) Prolonged tunicamycin treatment leads to the suppression of the mTORC1 pathway. Western blot assessment of mTORC1 activity in HeLa cells treated with 1 µg/ml tunicamycin for the indicated timepoints. 8 hours of treatment with tunicamycin leads to substantial inhibition of mTORC1, assessed by reduced levels of p-p70S6K and p-RPS6.

**Supplementary Figure 2 Support to main Figure 1**. (**A-B**) Metagene profiles of footprints aligned to either the start codon (A) or the stop codon (B) of all main Open Reading Frames, for DMSO (vehicle negative control) or tunicamycin-treated HeLa cells.

(A) Ribosome profiling on HeLa cells treated for 16-hours with tunicamycin reveals global translational and transcriptional reprogramming. Scatter plot showing change in translation (x-axis) versus change in mRNA levels (y-axis). This prolonged treatment causes induction of transcripts that affect the mTORC1 pathway - SESN2 and DDIT4.

(A) uORFs govern both ISR-mediated induction (ATF4) and suppression (ODC1) of the reporter’s translation. Mutation of the uORF start codon AUG to UAC abolishes these effects. n=3 Significance by unpaired, two-sided, t-test and adjusted for multiple testing. Error bars: standard deviation. p-values: * p<0.05, ** p<0.01, *** p < 0.001, ns = not significant.

**Supplementary Figure 3 Presence of either a uORF or an oORF is sufficient to cause induction of a ZBTB45 reporter. (A)** Schematic diagram illustrating the canonical delayed-reinitiation mechanism thought to induce ATF4 expression in response to stress.

**(B)** A luciferase reporter carrying the 5’UTR of ZBTB45 is induced in response to stress as long as either the uORFs or the oORF are present, but both are not required.

The left panel shows relative expression levels of the reporters in non-stressed conditions. The right panel shows induction of the same reporters upon treatment with tunicamycin. In all cases, uORF or oORF start codons were mutated to UAC. n=3 Significance by unpaired, two-sided, t-test and adjusted for multiple testing. Error bars: standard deviation. p-values: * p<0.05, ** p<0.01, *** p < 0.001, ns = not significant.

**Supplementary Figure 4** (A) Translational and transcriptional response to 2h of tunicamycin for the genes from which 5’UTRs were cloned to make the reporters shown in main Figure 2A. Each transcript is predicted to have only one overlapping ORF and no uORF.

(B) The DCP2 reporter is induced in response to the integrated stress response, but not mTORC1 inhibition. Treatment with 250nM Torin, which does not affect eIF2a phosphorylation, does not cause induction of the DCP2 reporter. n=3 Significance by unpaired, two-sided, t-test and adjusted for multiple testing. Error bars: standard deviation. p-values: * p<0.05, ** p<0.01, *** p < 0.001, ns = not significant.

**Supplementary Figure 5 Stress causes an equal reduction in translation of a library of reporters with different Kozak sequences.** (A) Schematic diagram illustrating the mechanistic hypothesis being tested in this figure. One possibility is that activation of the Integrated Stress Response causes translation initiation to drop by differing degrees depending on the identity of the Kozak sequence. If stress causes initiation to drop more strongly on the oORF start codon than the mORF start codon, this would lead to a relative induction in mORF translation.

(B) A library of bicistronic reporters was constructed whereby Renilla Luciferase (RLuc) has an oORF with Kozak sequences of varying strengths and Firefly Luciferase (FLuc) has the LMNB1 control 5’UTR lacking any uORFs or oORFs. Fluc and Rluc are driven by two independent CMV promoters. All transfections were normalised to a control reporter in which both Rluc and Fluc were expressed from uORF-less LMNB1 5′ UTRs.

(C) Scatter plot showing the Rluc/Fluc ratio (y-axis), reflecting the extent of Rluc suppression imposed by each cloned Kozak sequence, plotted against the estimated oORF Kozak strength obtained from (Noderer *et al.*, 2014).

(D) Tunicamycin treatment does not alter the relative strength of different Kozak sequences. Each point represents one reporter carrying RLuc with an oORF with a specific Kozak sequence. Relative RLuc activity in the control, non-stressed condition is plotted on the x-axis, and RLuc activity upon ISR induction on the y-axis. The linear regression line (black) is very similar to the y=x diagonal (red) with no data points significantly deviating from the regression, indicating that all Kozak sequences are equally repressed upon ISR induction. Significance by multiple unpaired t-test. n=5, ns=not significant. Error bars = standard deviation.

**Supplementary Figure 6 A hairpin in the DCP2 oORF is not needed for inducibility of the DCP2 reporter.** (A) Distribution of Kozak strengths for uORFs of ISR-induced transcripts, overlaid on the histogram representing the strength distribution of all Kozak sequences previously tested in (Noderer *et al.*, 2014).

(B) Reporter expression in basal, non-stressed conditions for the reporters shown in main Figure 3C.

(C) Structure of the hairpin present in the DCP2 oORF, predicted by Forna (Kerpedjiev *et al*, 2015). Mutations, used for the two constructs, aimed to disrupt the stem of the structure, are indicated. The amino acid sequence of the encoded peptide is indicated in one-letter code.

(D) Modification of the DCP2 oORF hairpin does not affect DCP2-reporter inducibility. n=3 Significance by unpaired, two-sided, t-test and adjusted for multiple testing. Error bars: standard deviation. p-values: * p<0.05, ** p<0.01, *** p < 0.001, ns = not significant.

**Supplementary Figure 7 Inducibility of the DCP2 reporter depends on a repressive element at the 3’ end of the oORF.** (A) Detailed sequences of the chimeric oORFs created by combining the DCP2 oORF with either a synthetic oORF or the RNFT1 oORF, assayed in main Figure 4A-B. Parts derived from each reporter are color coded.

(B) Schematic representation and detailed sequence of the premature stop codon introduced into the DCP2 oORF.

(C) Truncation of the DCP2 oORF causes an increase in the basal expression of the DCP2 reporter.

(D) The C-terminus of the DCP2 oORF peptide is highly conserved across vertebrate species

(E) Detailed amino acid sequences resulting from mutations introduced into the DCP2 oORF.

**Supplementary Figure 8 Expression of the DCP2 oORF increases in response to ISR activation.** (A) Schematic diagram illustrating two different mechanistic hypotheses how the oORF 3’ end could cause induction of mORF translation in response to stress. In one case (top), it affects leaky scanning through the oORF start site, in which case translation of the oORF should decrease upon ISR activation. In the other case (bottom), the oORF 3’end acts as a translation block for both the oORF and the mORF, which is relieved by ISR activation. In this case, oORF translation should increase upon ISR activation.

(B) Tunicamycin treatment leads to the accumulation of HA-tagged DCP2-oORF peptide. ATF4 is used as a control for ISR induction, and tubulin as a loading control. Quantification of HA/tubulin ratio is shown on the right. n=3 Significance by unpaired, two-sided, t-test and adjusted for multiple testing. Error bars: standard deviation. p- values: ns = not significant.

(C) The DCP2 oORF strongly reduces expression of Rluc when fused in-frame. Expression levels in non-stressed cells are shown.

(D) RLuc fused in-frame to the DCP2 oORF increases in expression upon tunicamycin treatment. n=4 Significance by unpaired, two-sided, t-test and adjusted for multiple testing. Error bars: standard deviation. p-values: * p<0.05, *** p < 0.001, ns = not significant.

**Supplementary Figure 9 Support to main Figure 5** (A) The DCP2 oORF causes induction of a luciferase reporter upon ISR activation also when placed as a uORF within a heterologous 5’UTR (LMNB1). The DCP2 main ORF start codon, which is present inside the DCP2 oORF CDS, was mutated to UAG to prevent initiation in a separate coding frame. n=3 Significance by unpaired, two-sided, t-test and adjusted for multiple testing. Error bars: standard deviation. p-values: ** p<0.01, ns = not significant.

(B) Reporter expression in basal, non-stressed conditions for the reporters shown in main Figure 5B.

(C) Detailed amino acid sequences resulting from mutations introduced into the ATF- 4 oORF. Neither of the mutations changes the ATF4 main ORF.

## References

1. Adomavicius T, Guaita M, Zhou Y, Jennings MD, Latif Z, Roseman AM, Pavitt GD (2019) The structural basis of translational control by eIF2 phosphorylation. Nature communications 10: 2136

2. Andreev DE, O’Connor PB, Fahey C, Kenny EM, Terenin IM, Dmitriev SE, Cormican P, Morris DW, Shatsky IN, Baranov PV (2015) Translation of 5’ leaders is pervasive in genes resistant to eIF2 repression. eLife 4: e03971

3. Bogorad AM, Lin KY, Marintchev A (2017) Novel mechanisms of eIF2B action and regulation by eIF2alpha phosphorylation. Nucleic Acids Res 45: 11962–11979

4. Bohlen J, Roiuk M, Neff M, Teleman AA (2023) PRRC2 proteins impact translation initiation by promoting leaky scanning. Nucleic Acids Res 51: 3391–3409

5. Condon KJ, Orozco JM, Adelmann CH, Spinelli JB, van der Helm PW, Roberts JM, Kunchok T, Sabatini DM (2021) Genome-wide CRISPR screens reveal multitiered mechanisms through which mTORC1 senses mitochondrial dysfunction. Proceedings of the National Academy of Sciences of the United States of America 118

6. Costa-Mattioli M, Walter P (2020) The integrated stress response: From mechanism to disease. Science 368

7. Dever TE, Ivanov IP, Hinnebusch AG (2023) Translational regulation by uORFs and start codon selection stringency. Genes Dev 37: 474–489

8. Figlia G, Muller S, Garcia-Cortizo F, Neff M, Klinke G, Poschet G, Teleman AA (2025) mTORC1 senses glutamine and other amino acids through GCN2. The EMBO journal 44: 4825–4866

9. Han P, Shichino Y, Schneider-Poetsch T, Mito M, Hashimoto S, Udagawa T, Kohno K, Yoshida M, Mishima Y, Inada T et al (2026) Genome-wide Survey of Ribosome Collision. Cell reports 45: 116845

10. Harding HP, Novoa I, Zhang Y, Zeng H, Wek R, Schapira M, Ron D (2000) Regulated translation initiation controls stress-induced gene expression in mammalian cells. Mol Cell 6: 1099–1108

11. Hashimoto Y, Zhang C, Kawauchi J, Imoto I, Adachi MT, Inazawa J, Amagasa T, Hai T, Kitajima S (2002) An alternatively spliced isoform of transcriptional repressor ATF3 and its induction by stress stimuli. Nucleic Acids Res 30: 2398–2406

12. Heberle AM, Prentzell MT, van Eunen K, Bakker BM, Grellscheid SN, Thedieck K (2015) Molecular mechanisms of mTOR regulation by stress. Mol Cell Oncol 2: e970489

13. Heldens L, Hensen SM, Onnekink C, van Genesen ST, Dirks RP, Lubsen NH (2011) An atypical unfolded protein response in heat shocked cells. PloS one 6: e23512

14. Hetz C, Zhang K, Kaufman RJ (2020) Mechanisms, regulation and functions of the unfolded protein response. Nature reviews Molecular cell biology 21: 421–438

15. Ho DV, Chan JY (2015) Induction of Herpud1 expression by ER stress is regulated by Nrf1. FEBS letters 589: 615–620

16. Johnstone TG, Bazzini AA, Giraldez AJ (2016) Upstream ORFs are prevalent translational repressors in vertebrates. The EMBO journal 35: 706–723

17. Kashiwagi K, Yokoyama T, Nishimoto M, Takahashi M, Sakamoto A, Yonemochi M, Shirouzu M, Ito T (2019) Structural basis for eIF2B inhibition in integrated stress response. Science 364: 495–499

18. Kenner LR, Anand AA, Nguyen HC, Myasnikov AG, Klose CJ, McGeever LA, Tsai JC, Miller-Vedam LE, Walter P, Frost A (2019) eIF2B-catalyzed nucleotide exchange and phosphoregulation by the integrated stress response. Science 364: 491–495

19. Kerpedjiev P, Hammer S, Hofacker IL (2015) Forna (force-directed RNA): Simple and effective online RNA secondary structure diagrams. Bioinformatics 31: 3377–3379

20. Kwon OS, An S, Kim E, Yu J, Hong KY, Lee JS, Jang SK (2017) An mRNA-specific tRNAi carrier eIF2A plays a pivotal role in cell proliferation under stress conditions: stress-resistant translation of c-Src mRNA is mediated by eIF2A. Nucleic Acids Res 45: 296–310

21. Lu PD, Harding HP, Ron D (2004) Translation reinitiation at alternative open reading frames regulates gene expression in an integrated stress response. The Journal of cell biology 167: 27–33

22. Noderer WL, Flockhart RJ, Bhaduri A, Diaz de Arce AJ, Zhang J, Khavari PA, Wang CL (2014) Quantitative analysis of mammalian translation initiation sites by FACS- seq. Molecular systems biology 10: 748

23. Pakos-Zebrucka K, Koryga I, Mnich K, Ljujic M, Samali A, Gorman AM (2016) The integrated stress response. EMBO reports 17: 1374–1395

24. Pestova TV, Kolupaeva VG (2002) The roles of individual eukaryotic translation initiation factors in ribosomal scanning and initiation codon selection. Genes Dev 16: 2906–2922

25. Proud CG (2019) Phosphorylation and Signal Transduction Pathways in Translational Control. Cold Spring Harbor perspectives in biology 11

26. Reich S, Nguyen CDL, Has C, Steltgens S, Soni H, Coman C, Freyberg M, Bichler A, Seifert N, Conrad D et al (2020) A multi-omics analysis reveals the unfolded protein response regulon and stress-induced resistance to folate-based antimetabolites. Nature communications 11: 2936

27. Roiuk M, Neff M, Teleman AA (2024) eIF4E-independent translation is largely eIF3d- dependent. Nature communications 15: 6692

28. Roiuk M, Neff M, Teleman AA (2025) Human eIF2A has a minimal role in translation initiation and in uORF-mediated translational control in HeLa cells. eLife 14

29. Schleich S, Acevedo JM, Clemm von Hohenberg K, Teleman AA (2017) Identification of transcripts with short stuORFs as targets for DENR*MCTS1-dependent translation in human cells. Scientific reports 7: 3722

30. Schleich S, Strassburger K, Janiesch PC, Koledachkina T, Miller KK, Haneke K, Cheng YS, Kuchler K, Stoecklin G, Duncan KE et al (2014) DENR-MCT-1 promotes translation re-initiation downstream of uORFs to control tissue growth. Nature 512: 208–212

31. Sidrauski C, McGeachy AM, Ingolia NT, Walter P (2015) The small molecule ISRIB reverses the effects of eIF2alpha phosphorylation on translation and stress granule assembly. eLife 4

32. Skabkin MA, Skabkina OV, Hellen CU, Pestova TV (2013) Reinitiation and Other Unconventional Posttermination Events during Eukaryotic Translation. Mol Cell

33. Smirnova AM, Hronova V, Mohammad MP, Herrmannova A, Gunisova S, Petrackova D, Halada P, Coufal S, Swirski M, Rendleman J et al (2024) Stem-loop-induced ribosome queuing in the uORF2/ATF4 overlap fine-tunes stress-induced human ATF4 translational control. Cell reports 43: 113976

34. Smirnova VV, Shestakova ED, Nogina DS, Mishchenko PA, Prikazchikova TA, Zatsepin TS, Kulakovskiy IV, Shatsky IN, Terenin IM (2022) Ribosomal leaky scanning through a translated uORF requires eIF4G2. Nucleic Acids Res 50: 1111–1127

35. Starck SR, Tsai JC, Chen K, Shodiya M, Wang L, Yahiro K, Martins-Green M, Shastri N, Walter P (2016) Translation from the 5’ untranslated region shapes the integrated stress response. Science 351: aad3867

36. Vattem KM, Wek RC (2004) Reinitiation involving upstream ORFs regulates ATF4 mRNA translation in mammalian cells. Proceedings of the National Academy of Sciences of the United States of America 101: 11269–11274

37. Wang XQ, Rothnagel JA (2004) 5’-untranslated regions with multiple upstream AUG codons can support low-level translation via leaky scanning and reinitiation. Nucleic Acids Res 32: 1382–1391

38. Weiss B, Dikstein R (2024) Unraveling the landscapes and regulation of scanning, leaky scanning, and 48S initiation complex conformations. Cell reports 43: 114126

39. Wek RC, Staschke KA (2010) How do tumours adapt to nutrient stress? The EMBO journal 29: 1946–1947

40. Whitney ML, Jefferson LS, Kimball SR (2009) ATF4 is necessary and sufficient for ER stress-induced upregulation of REDD1 expression. Biochemical and biophysical research communications 379: 451–455

41. Ye J, Palm W, Peng M, King B, Lindsten T, Li MO, Koumenis C, Thompson CB (2015) GCN2 sustains mTORC1 suppression upon amino acid deprivation by inducing Sestrin2. Genes Dev 29: 2331–2336

42. Yoshida H, Matsui T, Yamamoto A, Okada T, Mori K (2001) XBP1 mRNA is induced by ATF6 and spliced by IRE1 in response to ER stress to produce a highly active transcription factor. Cell 107: 881–891

43. Young SK, Palam LR, Wu C, Sachs MS, Wek RC (2016) Ribosome Elongation Stall Directs Gene-specific Translation in the Integrated Stress Response. The Journal of biological chemistry 291: 6546–6558

44. Young SK, Wek RC (2016) Upstream Open Reading Frames Differentially Regulate Gene-specific Translation in the Integrated Stress Response. The Journal of biological chemistry 291: 16927–16935

45. Young SK, Willy JA, Wu C, Sachs MS, Wek RC (2015) Ribosome Reinitiation Directs Gene-specific Translation and Regulates the Integrated Stress Response. The Journal of biological chemistry 290: 28257–28271

46. Zhou J, Wan J, Shu XE, Mao Y, Liu XM, Yuan X, Zhang X, Hess ME, Bruning JC, Qian SB (2018) N(6)-Methyladenosine Guides mRNA Alternative Translation during Integrated Stress Response. Mol Cell 69: 636–647 e637

