## Supplementary Figures for "Stress induces DCP2 translation via a stalling-dependent mechanism"

### Suppl. Figure 1

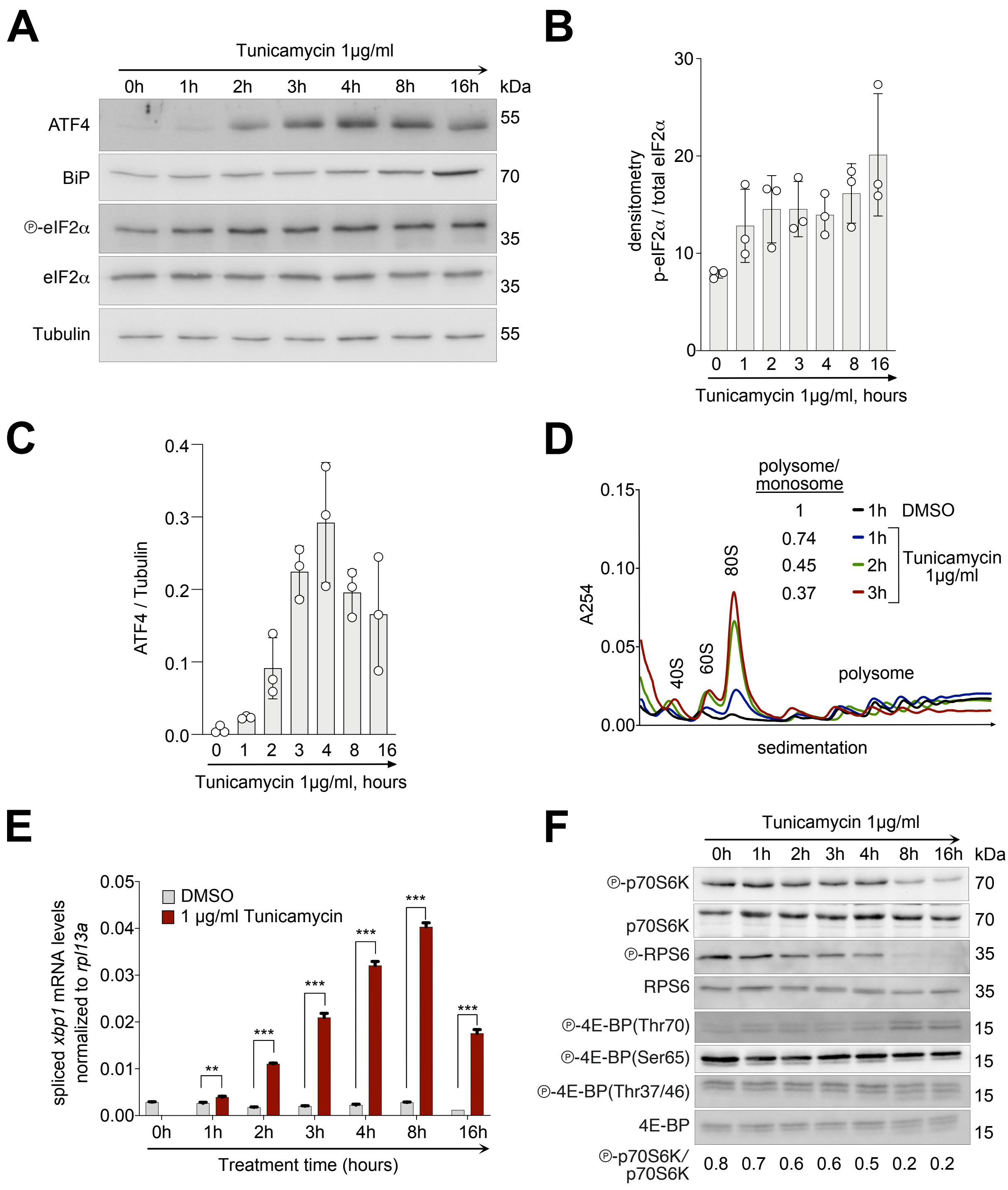

### Suppl. Figure 2

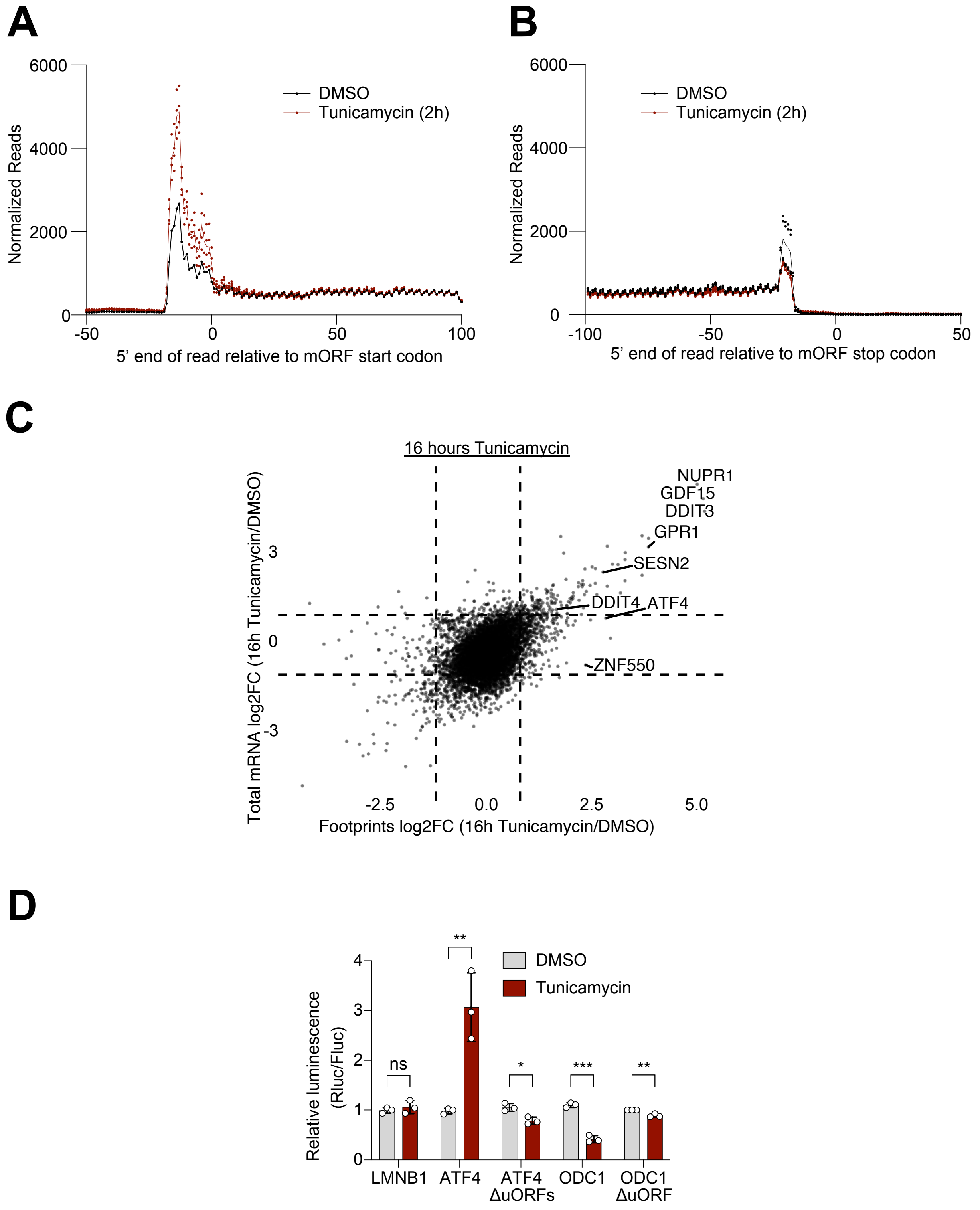

### Suppl. Figure 3

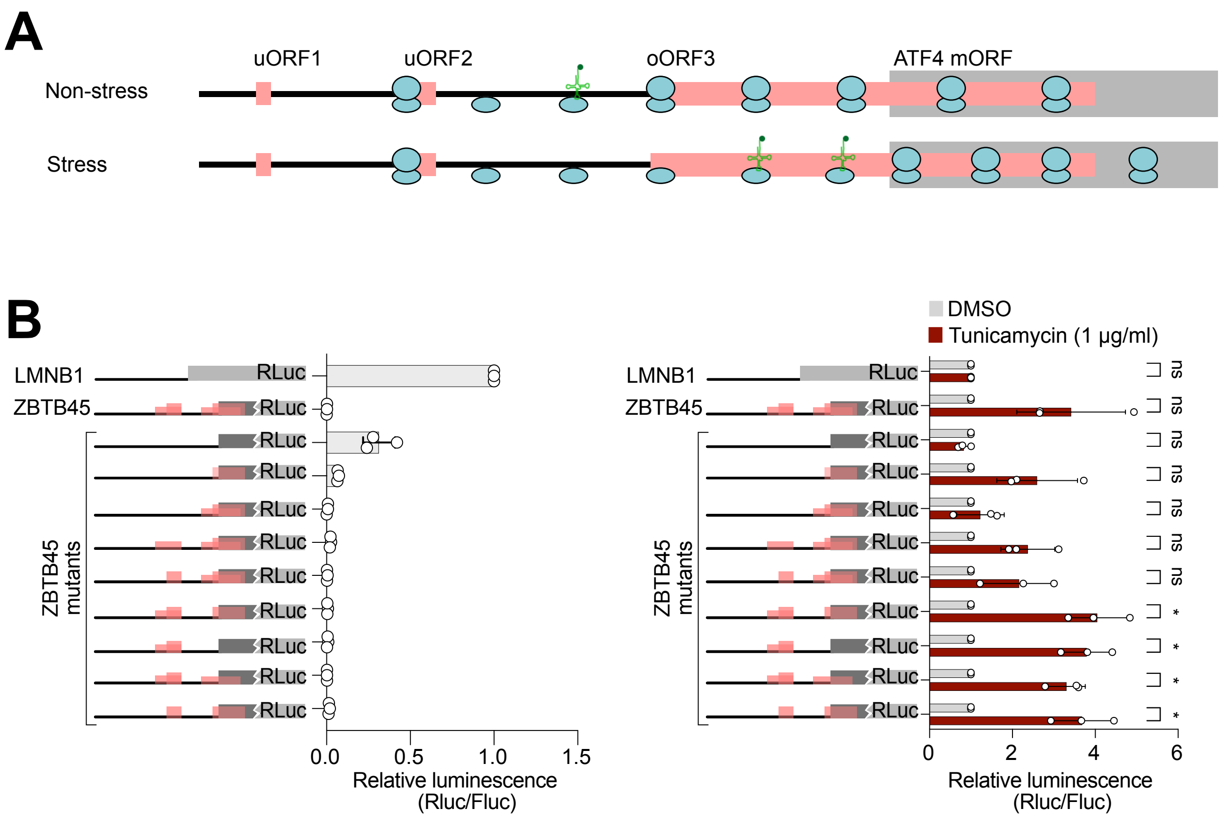

### Suppl. Figure 4

A

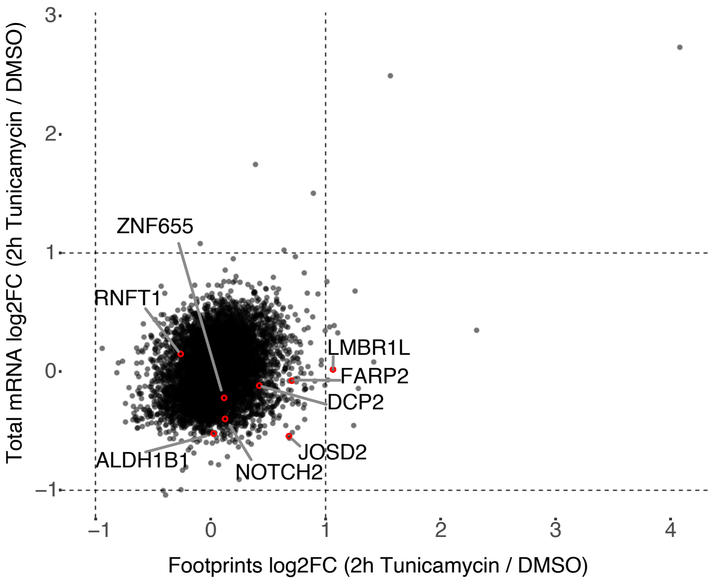


B

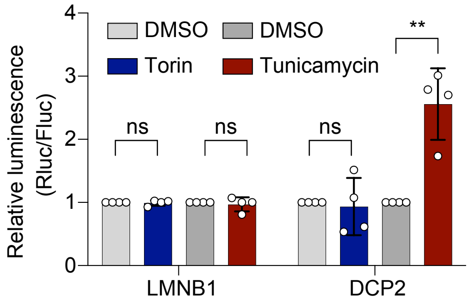

### Suppl. Figure 5

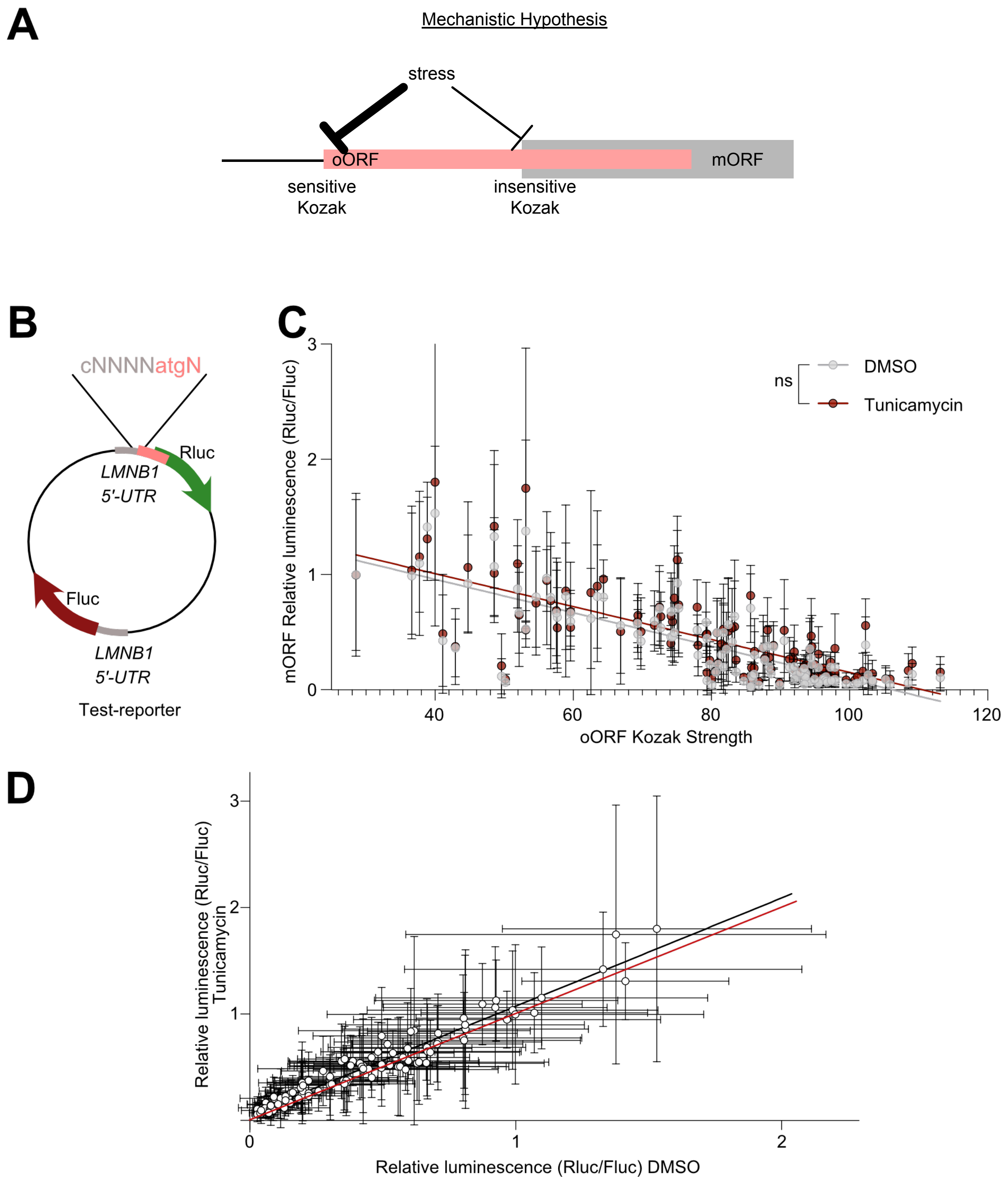

### Suppl. Figure 6

A

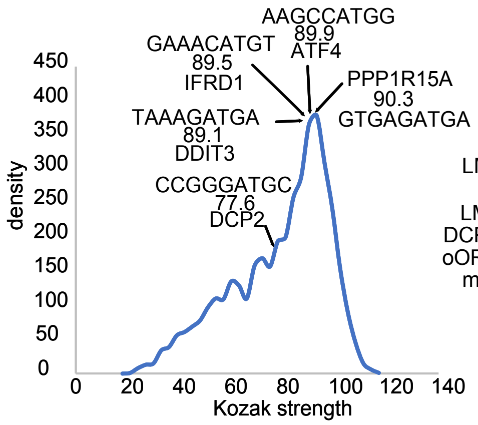


B

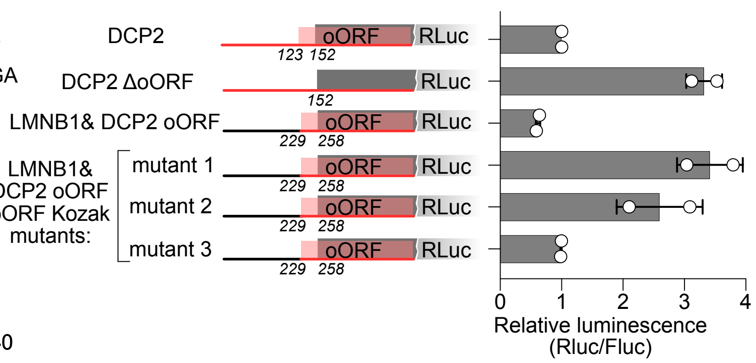


C

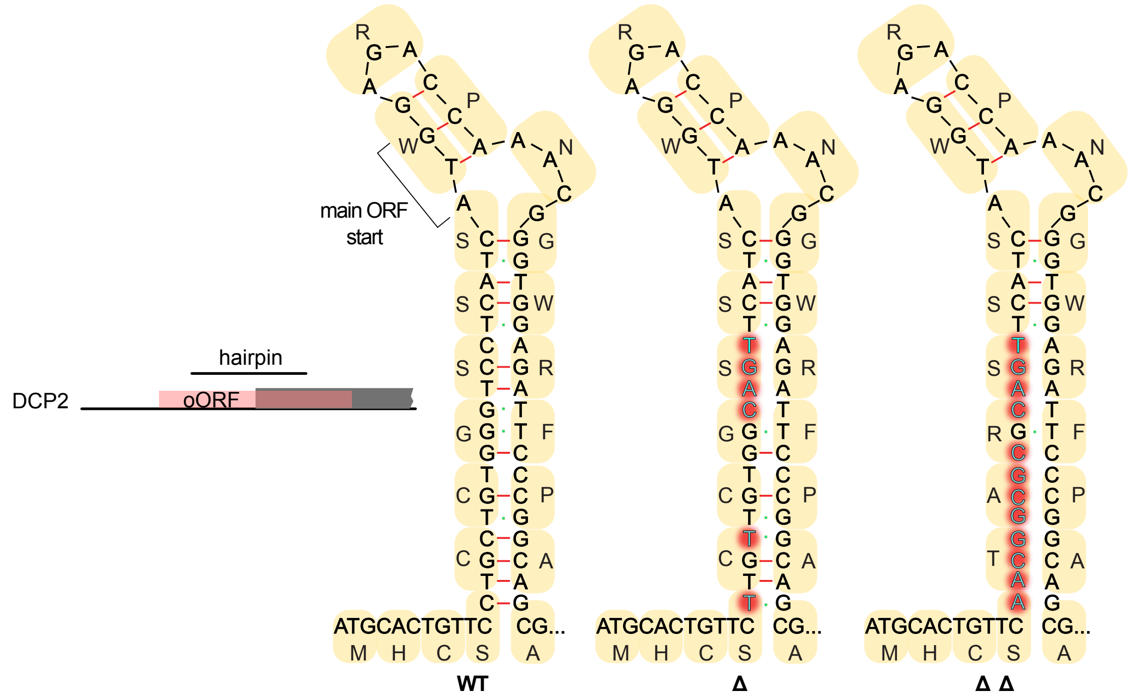


D

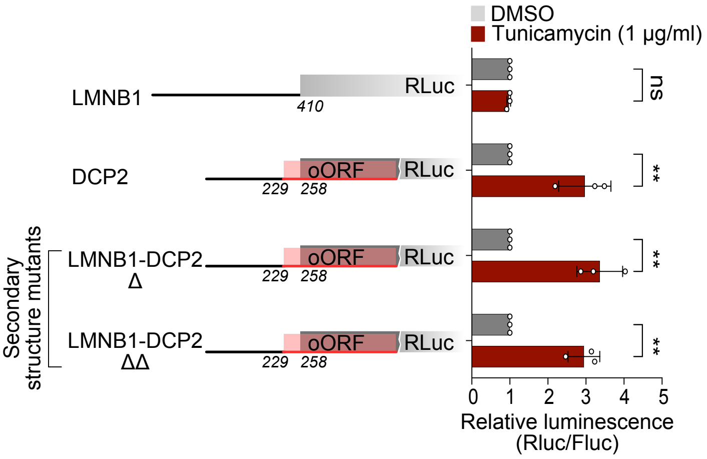

### Suppl. Figure 7

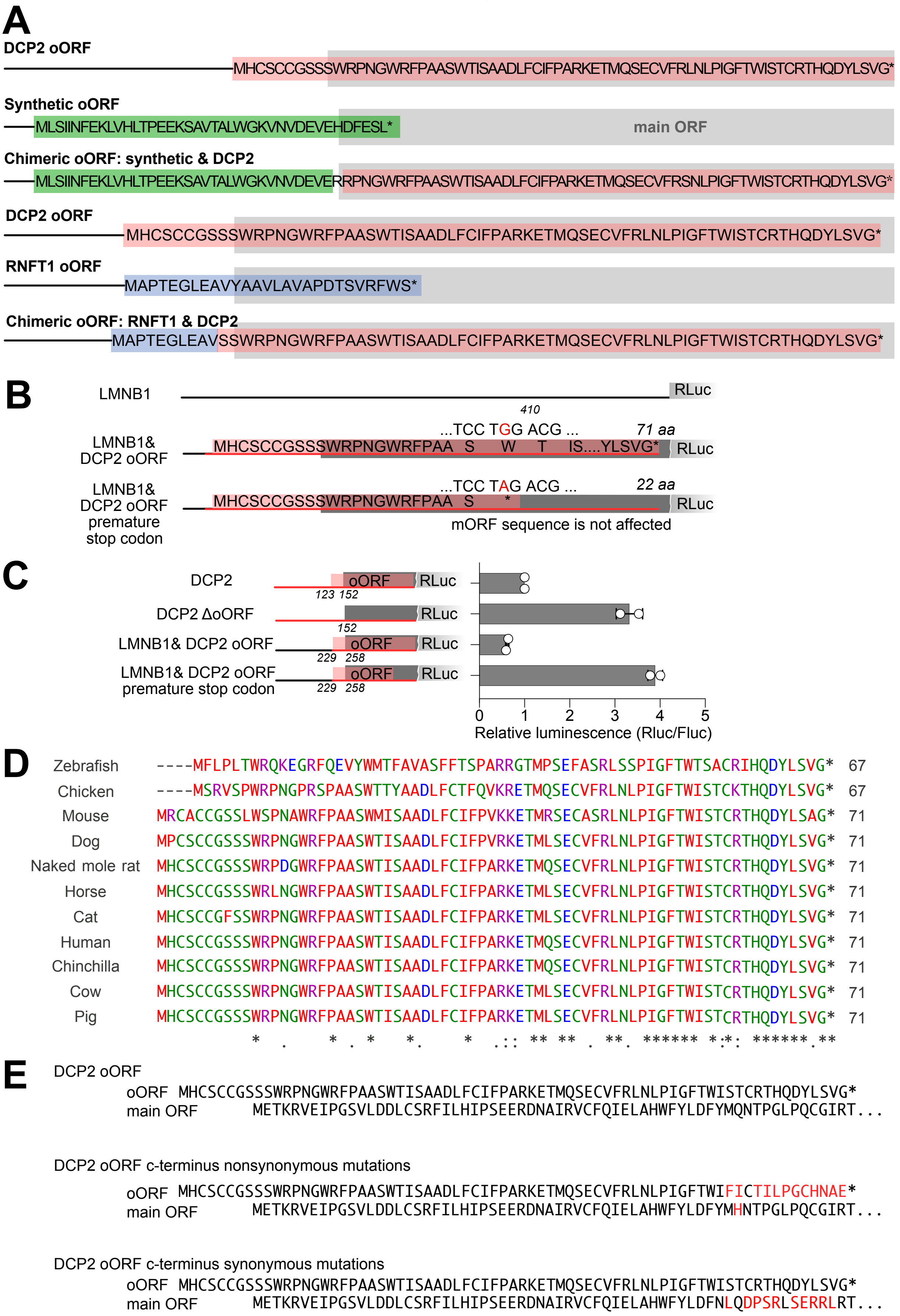

### Suppl. Figure 8

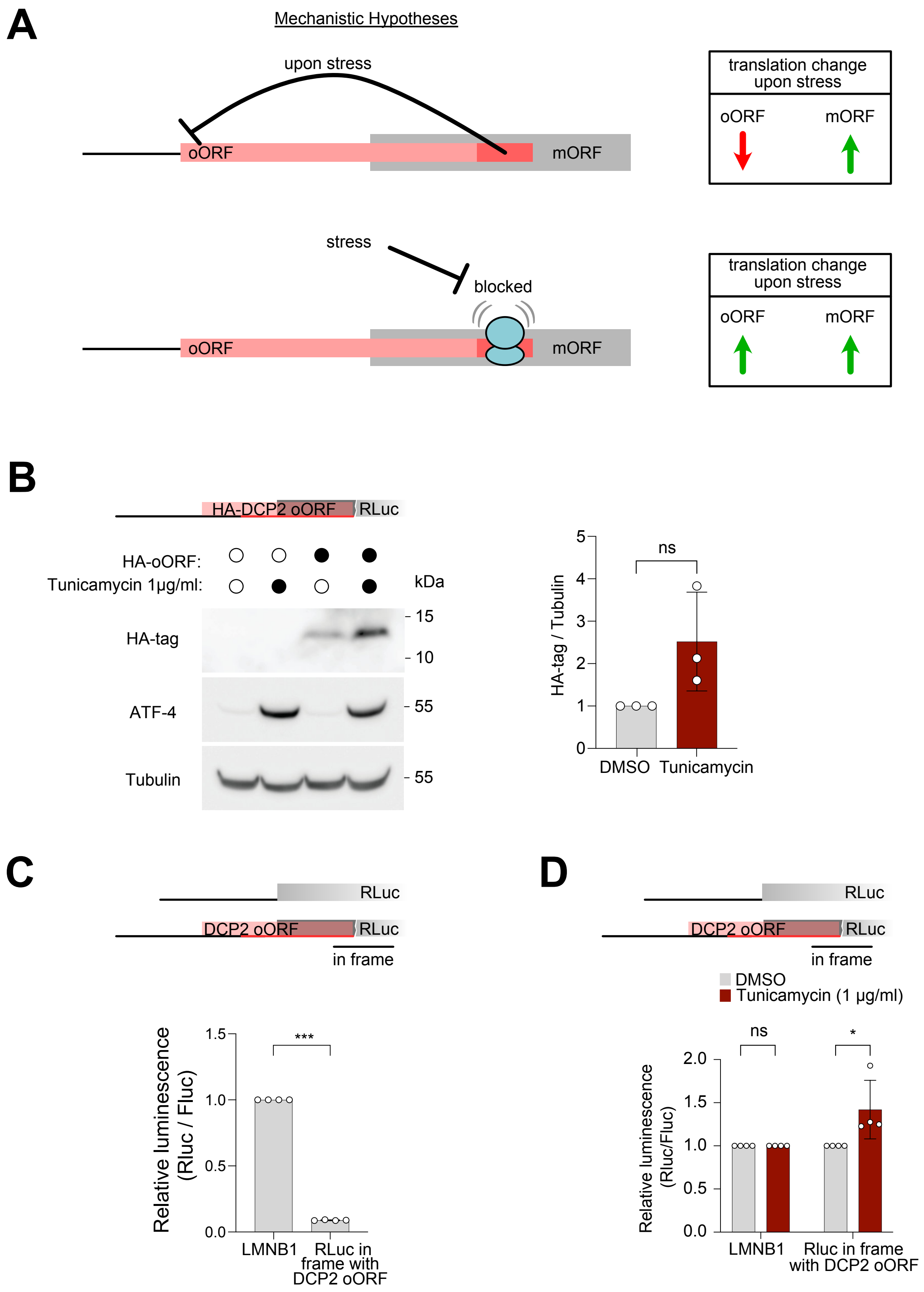

### Suppl. Figure 9

## A

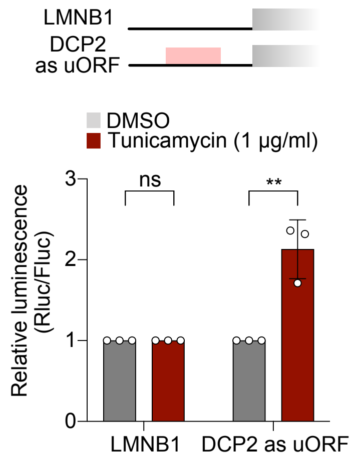


## B

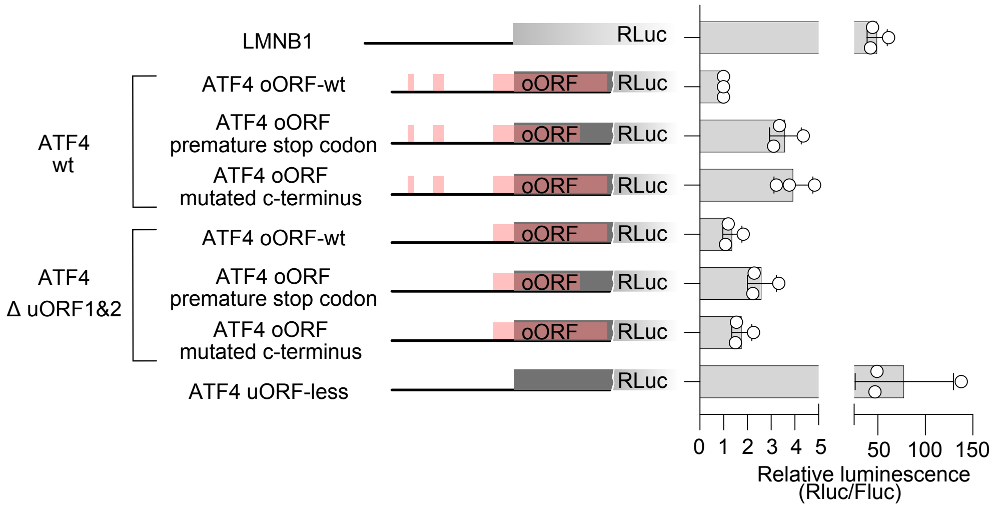


## C

ATF4 oORF WT

oORF MALLTAFSSSVAVTDKDTFELSTFLDSSKAPQHDRNELPEQRGVGGGLDVPLRPVGFGG*
main ORF MTEMSFLSSEVLVGDLMSPFDQSGLGAES...

ATF4 oORF premature stop

oORF MALLTAFSSSVAVTDKDTFELSTFLDSSKAPQHDRNELPEQ*
main ORF MTEMSFLSSEVLVGDLMSPFDQSGLGAES...

ATF4 oORF mutated C-terminus

oORF MALLTAFSSSVAVTDKDTFELSTFLDSSKAPQHDRNELPEQRGVSRRFNVTVRSIRLRS*
main ORF MTEMSFLSSEVLVGDLMSPFDQSGLGAES...
